# The effects of epistasis and linkage on the invasion of locally beneficial mutations and the evolution of genomic islands

**DOI:** 10.1101/2021.04.30.442097

**Authors:** Martin Pontz, Reinhard Bürger

**Affiliations:** School of Zoology, Faculty of Life Sciences, Tel Aviv University,Israel; Department of Mathematics, University of Vienna, Austria; Vienna Graduate School of Population Genetics

**Keywords:** Local adaptation, Selection, Recombination, Invasion Probability, Gene Flow, Genetic Interactions

## Abstract

We study local adaptation of a peripheral population by investigating the fate of new mutations using a haploid two-locus two-allele continent-island migration model. We explore how linkage, epistasis, and maladaptive gene flow affect the invasion probability of weakly beneficial de-novo mutations that arise on the island at an arbitrary physical distance to a locus that already maintains a stable migration-selection polymorphism. By assuming a slightly supercritical branching process, we deduce explicit conditions on the parameters that permit a positive invasion probability and we derive approximations for it. They show how the invasion probability depends on the additive and epistatic effects of the mutant, on its linkage to the polymorphism, and on the migration rate. We use these approximations together with empirically motivated distributions of epistatic effects to analyze the influence of epistasis on the expected invasion probability if mutants are drawn randomly from such a distribution and occur at a random physical distance to the existing polymorphism. We find that the invasion probability generally increases as the epistasis parameter increases or the migration rate decreases, but not necessarily as the recombination rate decreases. Finally, we shed light on the size of emerging genomic islands of divergence by exploring the size of the chromosomal neighborhood of the already established polymorphism in which 50% or 90% of the successfully invading mutations become established. These ‘window sizes’ always decrease in a reverse sigmoidal way with stronger migration and typically increase with increasing epistatic effect.

## 1 Introduction

Local adaptation of a population to a new environment is frequently considered as a first step in a process leading to the emergence of a new species, because it increases the divergence between spatially separated populations. Often, gene flow will counteract this adaptive divergence. Local adaptation may occur from standing genetic variation or from new mutations. Gene flow is especially detrimental for a peripheral population if new, locally beneficial mutations need to be established to improve adaptation. Since a de-novo mutation starts out as a single copy, it is particularly prone to rapid loss caused by random events. As already shown by Haldane (1927) and Fisher (1930), the probability of survival of a mutant with selective advantage *s* is approximately 2*s*. Maladaptive gene flow reduces or even annihilates this probability.

We study the influence of linkage and epistasis on the fate of a new, weakly beneficial mutation in a peripheral (island) population that is exposed to maladaptive gene flow from the main (continental) population. For this purpose, we extend a branching process model of Aeschbacher and Bürger (2014) by accounting for genetic interactions between loci. By a well known dichotomy, the mutant is either lost by random drift, which usually occurs rather quickly, or it invades and becomes established in the population. In this case, its further growth trajectory is (primarily) determined by selection in concert with other deterministic forces. Beneficial mutations of large effect are believed to be very rare. If their selective advantage exceeds the immigration rate, they have a positive establishment probability unless they arise in tight linkage to a deleterious background. As in Aeschbacher and Bürger (2014), we assume that at some locus there already exists a stable polymorphism in the island population, which was established because the selective advantage of the new (island) mutant at this locus exceeded the immigration rate of the ancestral (continental) allele. Thus, a first step in local adaptation has already been achieved.

We explore under which conditions such a polymorphic locus can act as a starting point for further adaptation by facilitating establishment of locally beneficial alleles in its genomic neighborhood. Because weakly beneficial mutations are more likely to occur than strongly beneficial mutations (e.g., Orr, 2010; Bataillon and Bailey, 2014; Rice et al., 2015), we focus on the case where the selective advantage of a new mutation on the island is smaller than that of the island mutant at the polymorphic locus. Then the genetic background in which the mutant occurs, and which contains the already established polymorphism, plays a crucial role in enabling invasion and survival of the mutant.

Previous studies (Aeschbacher and Bürger, 2014; Yeaman et al., 2016) investigated a similar question for diploids, however, with the restriction to a genic selection regime, i.e., by ignoring dominance and epistasis. They showed that the probability of establishment of the new mutation is always higher for loci that are tightly linked to the existing polymorphism than for loosely linked loci, although it is not always maximized for complete linkage. Their interpretation was that this process favors the emergence of genomic regions containing clusters of locally beneficial mutations (Yeaman and Whitlock, 2011), or at least of loci contributing to divergence. Such regions were dubbed genomic islands of divergence (Nosil et al., 2009; Feder and Nosil, 2010), of speciation (Turner et al., 2005), or of differentiation (Harr, 2006).

We extend and complement these results by taking into account genetic interactions between new locally beneficial mutations and the existing polymorphism. The invasion probability generally increases with more positive epistasis and mostly also with less recombination. However, it may be maximized at a positive recombination rate, especially if epistasis is positive and the additive effect not too small. Furthermore, the relative contributions of epistasis and linkage to the invasion probability depend crucially on the strength of gene flow. We find that for weak migration a change in the recombination rate has virtually no effect on the invasion probability, whereas an increase in epistasis has a clear positive effect. For moderate or high migration rates, however, tighter linkage elevates the invasion probability substantially, often more than an increase in epistasis.

Although we are in an era of relatively cheap sequencing technology and advanced bioinformatic tools, it is still unclear how epistatic values are distributed and which distribution is most prevalent in diploid organisms (e.g., Ehrenreich, 2017; Gao et al., 2010). More results are available for haploid organisms, because the fitness structure without dominance effects is simpler and also allows for mathematical approximations of the epistasis distribution (e.g., Martin et al., 2007; Blanquart et al., 2014; Schoustra et al., 2016). In order to take advantage of this fact and to limit mathematical complications, we investigate a haploid version of the general model.

Adaptation typically occurs on longer time scales, such that many new mutations may appear in a population, and their physical distance to the existing migrationselection polymorphism as well as their additive and epistatic effects on fitness may be considered as being drawn from appropriate distributions. Based on data-motivated distributions of the strength of epistasis, we especially analyze the expected effects of epistasis. By averaging the invasion probability of a mutation over such distributions, we can draw conclusions about the general importance of the involved evolutionary factors for the adaptation of a population and the expected genomic architecture. Concerning the latter, we follow Yeaman et al. (2016) and explore how the size of genomic islands of divergence will depend on migration rates, additive and epistastic effects, and their distributions.

## 2 Methods

### 2.1 Model and biological scenario

Our model is closely related to that employed by Aeschbacher and Bürger (2014), which is a diploid, discrete-time, two-locus two-allele model with continent-to-island (CI) migration. Whereas these authors assumed genic fitnesses, i.e., no dominance or epistasis, we include a parameter for genetic interaction between the two loci, but assume a haploid population.

The two loci are denoted by A and B and their alleles by *A*_1_, *A*_2_ and *B*_1_, *B*_2_. These form the four haplotypes *A*_1_*B*_1_, *A*_1_*B*_2_, *A*_2_*B*_1_, and *A*_2_*B*_2_, which occur at frequencies *x*_1_, *x*_2_, *x*_3_, and *x*_4_ on the island. We assume that the population on the continent is well adapted and fixed for alleles *A*_2_ and *B*_2_. The (im)migration rate to the island is denoted by *m*, i.e., each generation a fraction *m* of the adult population (after selection and recombination) on the island is replaced by individuals of the continental population. The recombination rate between locus A and locus B is denoted by *r*, and the allele frequencies of *A*_1_ and *B*_1_ on the island are *p* = *x*_1_ + *x*_2_ and *q* = *x*_1_ + *x*_3_, respectively.

Initially, the island population is fixed for *A*_2_, whereas at locus B the locally beneficial allele *B*_1_ has arisen some time ago and is in migration-selection balance, which requires that its selective advantage *b* exceeds the migration rate *m* (see below). Then a weakly beneficial mutation occurs at locus A, resulting in a single copy of *A*_1_. Its fate is determined by direct selection on locus A, linkage to the selected locus B, migration, genetic interaction between the loci, and random genetic drift.

We focus on the scenario, where the island population is so large that after an initial stochastic phase during which the mutant is either lost or increases to appreciable frequency, the dynamics becomes deterministic. This implies, under some technical conditions, that the fate of *A*_1_ is decided during the stochastic phase. If it survives this phase, it will reach an attractor in the interior of the state space (most likely, a fully polymorphic equilibrium), since no other boundary equilibrium attracts orbits from the interior (see Appendix A). This survival of the stochastic phase is what we synonymously call successful invasion or establishment.

### 2.2 Fitness and evolutionary dynamics

We use the following fitness scheme, which is general for a haploid two-locus two-allele model. The matrix 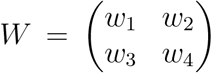 of relative genotypic fitnesses on the island is normalized such that

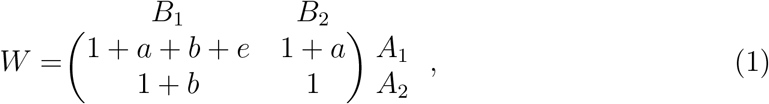

where *a* > 0 is the selective advantage of the new mutant *A*_1_ relative to the resident (continental) type *A*_2_ on the background *B*_2_; thus, *A*_1_*B*_2_ produces on average 1 + *a* times as many offspring as *A*_2_*B*_2_. Analogously, *b* > 0 is the selective advantage of *B*_1_ relative to *B*_2_ on the background *A*_2_. The parameter *e* measures epistasis, i.e., the deviation of the fitness of *A*_1_*B*_1_ from additivity. The only restriction we pose on *e* is −1 − *a* − *b < e*, so that the fitness of *A*_1_*B*_1_ is positive. We have negative (positive) epistasis if *e* < 0 (*e* > 0).

The evolutionary dynamics of such a haploid model depends on the life cycle of the population. We investigate only the dynamics in which selection occurs first, followed by recombination and migration. For weak evolutionary forces, the dynamics becomes independent of the order of selection, recombination, and migration (e.g. Bürger, 2014), and most explicit formulas simplify (e.g., Sect. 3.2).

Because selection occurs before recombination, we define the measure *D* of linkage disequilibrium after selection by

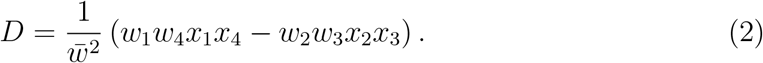

Writing *x_i_* and *x*′_*i*_ for the haplotpye frequencies in successive generations, the discrete-time dynamical equations can be expressed as

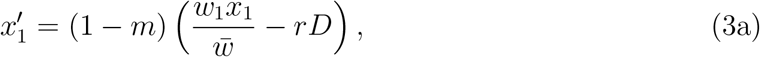

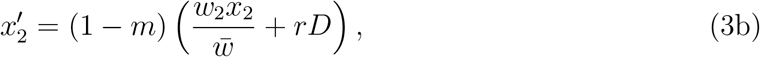

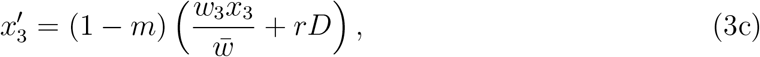

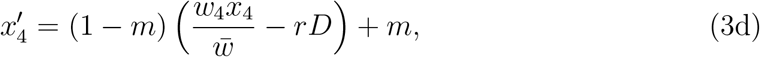

where the mean fitness is given by

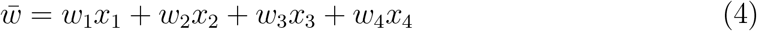

(cf. Felsenstein, 2019, chapter “Selection and Recombination”, p. 364).

Throughout this paper, we assume that the selective advantage *b* of *B*_1_ is large enough so that a stable polymorphism at B is maintained on the island population independently of any other locus. A brief calculation shows that this is case if and only if *b > m/*(1 − *m*) or, equivalently, if

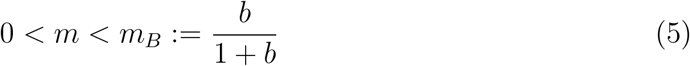

(cf. Haldane, 1930; Wright, 1931). Then the equilibrium frequency of *B*_1_ is

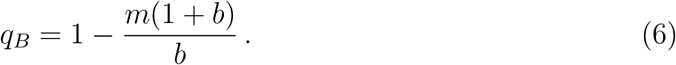

We call this equilibrium *E*_*B*_ and assume its existence. Therefore, a first step in local adaptation has been completed by establishing a polymorphism that persists despite maladaptive gene flow.

As argued above, we assume throughout that if a new beneficial mutant *A*_1_ occurs, it has a small fitness effect *a*, by which we mean *a < b*. We note that many of the tedious calculations performed to derive the results presented below are given in *Mathematica* notebooks (Wolfram Research 2020), which constitute the supplementary files S1 – S6.

Several formulas derived below are analogous to formulas derived in Aeschbacher and Bürger (2014) if epistasis is assumed absent. Nevertheless, even if we set *e* = 0, they differ algebraically in nontrivial ways because the model in Aeschbacher and Bürger (2014) is diploid, so that the new mutant *A*_1_ can occur on three genetic backgrounds, and they scaled the fitness of the double heterozygote to 1. Thus, already the equilibrium frequency *q*_*B*_ of *B*_1_ in our equation (6) differs from theirs. However, in the limit in which all evolutionary forces are weak, the mean matrices coincide if we set *e* = 0 (see Section 3.2).

### 2.3 Two-type branching process

We model the initial stochastic phase, after occurrence of *A*_1_, by a two-type branching process in discrete time (Harris, 1963). The two types are the haplotypes *A*_1_*B*_1_ and *A*_1_*B*_2_. Depending on the first occurrence of the mutant on background *B*_1_ or *B*_2_, the invasion probability of *A*_1_*B*_1_ or *A*_1_*B*_2_ is denoted by *π*_1_ or *π*_2_, respectively. The probability that *A*_1_ initially occurs in an individual with the *B*_1_ background depends on the equilibrium frequency of *B*_1_ at *E*_*B*_, which is *q*_*B*_ in (6).

The (mean) invasion probability 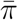 of *A*_1_ is thus the sum of the two conditional probabilities weighted by the frequencies of *B*_1_ and *B*_2_ at equilibrium:

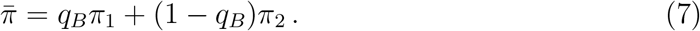

The invasion probability of *A*_1_ depends crucially on the so-called mean matrix **M** which, for a two-type process, is the 2 × 2 matrix with entries *λ*_*ij*_, where *λ*_*ij*_ is the mean number of *j*-type offspring produced by an *i*-type parent in each generation (while the mutant *A*_1_ is rare). The entries *λ*_*ij*_ can be obtained from of our basic recursion system (3) by computing the Jacobian *J* at the single-locus polymorphism *E*_*B*_ and identifying **M** as the transposed of the left upper 2 × 2 submatrix of *J*, which describes the dynamics of *A*_1_*B*_1_ and *A*_1_*B*_2_ (for details, see File S1, Sect. 6). Thus, the linearized dynamics around the equilibrium *E*_*B*_ pointing into the simplex is given by the recursion

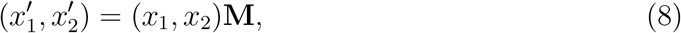

where, by a straightforward computation and then by substituting (1) into (9a),

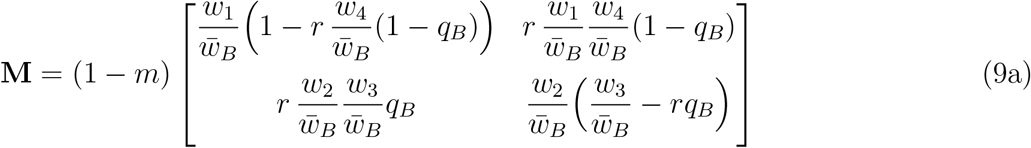

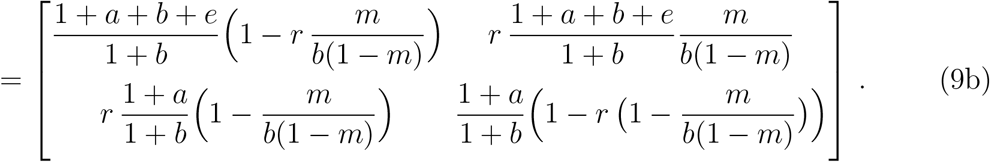

Here, 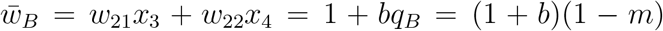 is the mean fitness at *E*_*B*_, and 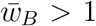 by (5). The off-diagonal terms in (9a) and (9b) are the mean number of offspring produced by recombination, i.e., the upper right (lower left) entry is the mean number of *A*_1_*B*_2_ (*A*_2_*B*_1_) offspring produced by *A*_1_*B*_1_ (*A*_2_*B*_2_) parents. The diagonal terms give the number of offspring produced without recombination. The sums of the top and bottom row, (1 + *a* + *b* + *e*)/(1 + *b*) and (1 + *a*)/(1 + *b*), measure the (total) reproductive ‘success’ of parents of type *A*_1_*B*_1_ and *A*_1_*B*_2_, respectively, when *A*_1_ is very rare and locus *B* is at the equilibrium *E*_*B*_.

It is well known (e.g. Harris, 1963) that invasion occurs with positive probability if and only if the leading eigenvalue *λ* of **M** satisfies

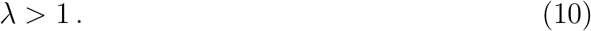

## 3 Analysis of the invasion condition

To apply the above theory to our model, we compute the leading eigenvalue *λ* of **M** and find (File S1, Sect. 6)

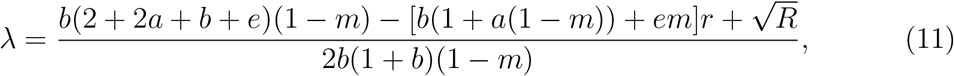

where

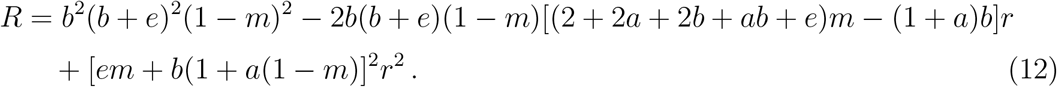

Using (11), we can rewrite the invasion condition (10) in the form

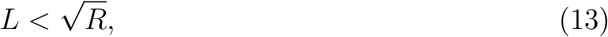

where

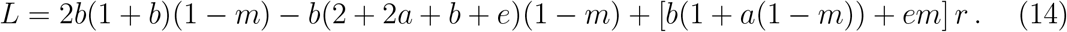

It will be useful to define *ϕ* = (*R* − *L*^2^)/[4*b*(1 − *m*)], which simplifies to

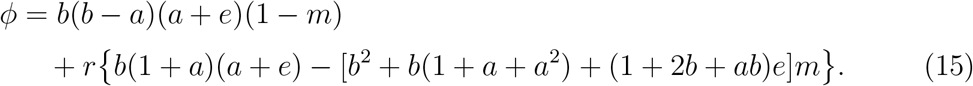

Note that 4*b*(1 − *m*) > 0 and thus does not influence the sign of *ϕ*, which is linear in both *m* and *r*. Therefore, the invasion condition (10) is satisfied if and only if *L* < 0 or *ϕ* > 0. In File S2, Sect. 2, we show that *L* < 0 implies *ϕ* > 0. As a consequence,

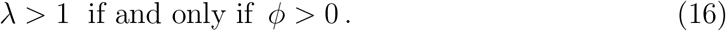

### 3.1 Characterization of the invasion condition in terms of the migration and the recombination rate

According to (16), invasion of *A*_1_ is possible if and only if *ϕ* > 0, where we recall our assumption 0 *< a < b*. Below, we define a critical migration rate *m*\* and then, alternatively, a critical recombination rate *r*\* to characterize the parameter region in which invasion of *A*_1_ can occur. The detailed derivations of the following results can be found in File S2, Sect. 3. We start by defining

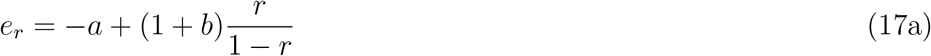

and

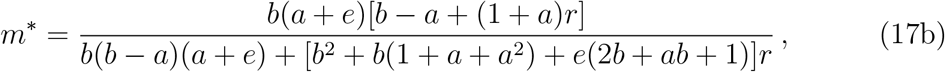

and note that *e*_*r*_ > 0 if and only if 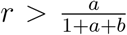, and *e < e*_*r*_ if and only if 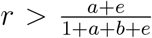. The following is our first main result.

#### Proposition 1.

i. *If* −1 − *a* − *b < e* ≤ −*a, then invasion is impossible.*
ii. *If −a < e < e*_*r*_, *then* 0 *< m*\* *< m*_*B*_ *and invasion is possible if m* ∈ [0, *m*\*).
iii. *If e*_*r*_ ≤ *e, then invasion is possible if m* ∈ [0, *m*_*B*_).

*Proof.* The proof can be simplified by transforming the linear function *ϕ*(*m*) in (15) into a function 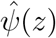 by substituting 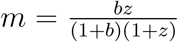. This strictly monotone transformation maps the interval 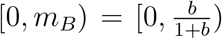 for admissible *m* (such that *E*_*B*_ exists) onto the interval [0, ∞) for *z*. Then 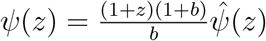 is linear in *z* and has the same sign as 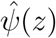. In fact, *ψ*(*z*) = *c*_0_ + *c*_1_*z*, where

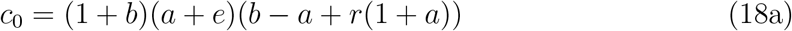

and

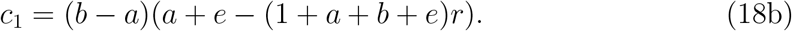

If −1 − *a* − *b < e* < −*a*, then *c*_0_ < 0 and *c*_1_ < 0, and thus *ψ*(*z*) < 0 if *z* > 0. This settles (i).

If −*a < e < e*_*r*_, then *c*_0_ > 0 and *c*_1_ < 0. Therefore, *ψ*(*z*) > 0 if *z* ∈ (0, *z*\*), where *z*\* is the zero of *ψ*(*z*). The corresponding zero of *ϕ*(*m*) is *m*\* given by (17b). In this case, 0 *< m*\* < *m*_*B*_ is clearly satisfied because by monotonicity of the transformation it is equivalent to 0 < *z*\* < ∞. Therefore, (ii) holds.

If *e > e*_*r*_, then *c*_0_ and *c*_1_ are positive. This implies that *ψ*(*z*) > 0 if *z* > 0, which in turn implies (iii).

As already noted above, *ϕ* is also a linear function in *r* and thus we can reformulate Proposition 1 in terms of *r*. We define

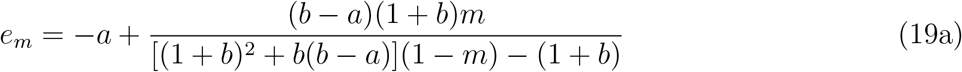

and

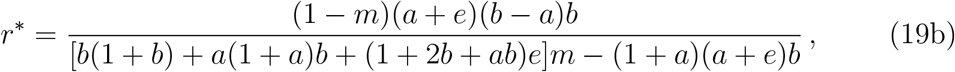

and note that *e*_*m*_ > 0 if *m > a*(1 − *a* + 2*b*)/(1 + *a* + *b* + 2*ab* − *a*^2^), where the lower bound is close to *a* if *a* and *b* are small. Now we can reformulate Proposition 1 as follows.

#### Proposition 2.

i. *If* −1 − *a* − *b < e* ≤ −*a, then invasion is impossible.*
ii. *If* −*a < e < e*_*m*_, *then* 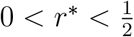 *and invasion is possible if r* ∈ [0, *r*\*).
iii. *If e*_*m*_ ≤ *e, then invasion is possible if* 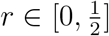.

*Proof.* With the transformation 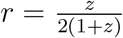, the proof is similar to that for Proposition 1.

Of course, the parameter regions described by Propositions 1 and 2 are identical. However, it is useful to have these two representations, because we want to investigate how epistasis interacts with linkage and with migration in admitting invasion of *A*_1_.

The following remark summarizes important properties of *λ*, *m*\*, and *r*\*.

#### Remark 3.

i. If *A* and *B* are completely linked (*r* = 0), then *M* and *λ* is independent of *m* and invasion of *A*_1_ is possible for every *m* if *a* + *e* > 0.
ii. The proof of Proposition 1 implies that *λ* = 1 if and only if *m* = *m*\* or, equivalently, *r* = *r*\*. Partially tedious computations, or applying *Mathematica’s* Reduce routine to the appropriate derivatives, show that (File S2, Sect. 3;)
iii. The eigenvalue *λ* has the following monotonicity properties. Every other parameter fixed, *λ* is strictly increasing in *a* and in *e*, and it is strictly decreasing in *m* and in *r*.
iv. If 0 < *m*\* < *m*_*B*_, then *m*\* is strictly increasing in *a* and in *e*, and *m*\* is strictly decreasing in *r*. In addition, *m*\* = 0 if *e* = −*a* and *m*\* = *m*_*B*_ if *e* = *e*_*r*_.
v. If 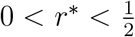, then *r*\* is strictly increasing in *a* and in *e*, and it is decreasing in *m*. In addition, *r*\* = 0 if *e* = −*a* and 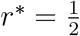 if *e* = *e*_*m*_.

The following remark summarizes immediate conclusions from Propositions 1 and 2 and Remark 3.

#### Remark 4.

i. Invasion of *A*_1_ is possible only if there is no sign epistasis, i.e., if *a*+*e* > 0, which means that the genotype *A*_1_*B*_1_ must have higher fitness in the fitness scheme (1) than any other genotype.
ii. Invasion of *A*_1_ is always facilitated by larger *a* and *e* and by smaller *m* and *r*. Moreover, *e*_*m*_ increases in *m* and *e*_*r*_ increases in *r*.
iii. For fixed selection coefficients *a*, *b*, and *e* satisfying *a* + *e* > 0, a reduction in *r* (*m*) will increase the maximum value of *m* (*r*) for which invasion is possible.

### 3.2 Weak-forces approximation

Several of the above expressions are complicated and do not easily provide analytical insight. However, if we assume that all evolutionary forces are weak, then much simpler expression are obtained. For this purpose, we scale the parameters for selection, migration, and recombination as

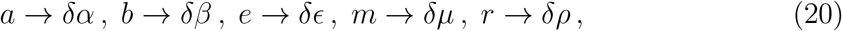

and assume that *δ* is small. By performing series expansions to first order in *δ* of the entries of the mean matrix **M** and returning to the original parameters (i.e., using roman font again), we obtain the weak-forces approximation

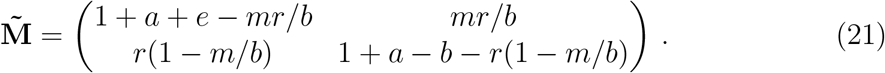

By applying the same procedure to every *x*′_*i*_ − *x*_*i*_ in the recursion system (3) and rescaling time according *τ* → *δt*, we obtain a system of approximating differential equations. Computing the Jacobian of this system at the equilibrium *E*_*B*_, which for this differential equation becomes *x*_1_ = *x*_2_ = 0, 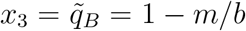, and *x*_4_ = *m/b*, we find that the upper left 2 × 2 submatrix is 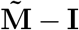, where **I** is the 2 × 2 identity matrix.

Throughout, we use the tilde ^~^ to denote the weak-forces approximation. The leading eigenvalue of 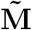 is

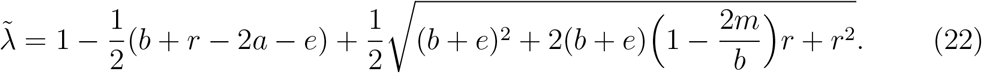

Series expansions to first order in *δ* using (20) show that the bounds used in Propositions 1 and 2 simplify to

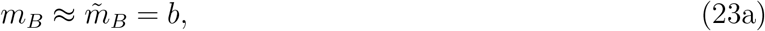

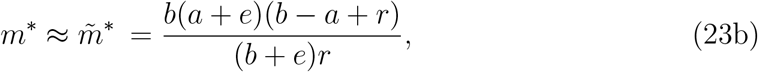

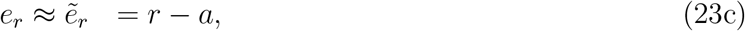

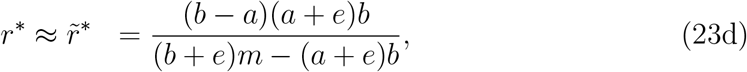

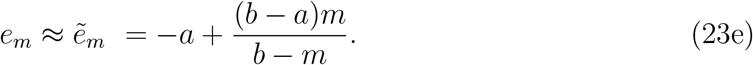

We note that if we set *e* = 0 in our mean matrix 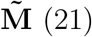, then it coincides with the mean matrix derived for weak evolutionary forces in Aeschbacher and Bürger (2014) (above eq. (53) in their supplementary File S1). Consequently, our leading eigenvalue 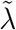 coincides with their *ν* (given above their eq. (55)), and our 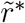 with their 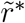 in (56).

With the above definitions, the following weak-forces versions of Propositions 1 and 2 are easily derived. They yield simple interpretations of the invasion conditions.

#### Proposition 5.

i. *If e* ≤ −*a, then invasion is impossible.*
ii. *If* −*a < e < r* − *a, then* 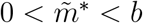 *and invasion is possible if 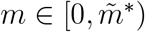.*
iii. *If r* − *a* ≤ *e, then invasion is possible if m* ∈ [0, *b*).

#### Proposition 6.

i. *If e* ≤ −*a, then invasion is impossible.*
ii. 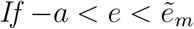, *then* 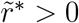 *and invasion is possible if* 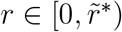.
iii. *If* 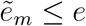, *then invasion is possible if r* ∈ [0, ∞).

Remarks 3 and 4 apply to these propositions as well. The computations for Sect. 3.2 are presented in File S5, Sects. 1,2.

## 4 Invasion probabilities

So far, we have characterized the parameter regions in which invasion of the new mutant *A*_1_ is possible, but we have not yet quantified the probability of invasion. In the following, we study the discrete-time branching process in which the number of offspring of type *j* produced by an individual of type *i* is Poisson distributed with mean *λ*_*ij*_, where the *λ*_*ij*_ are the entries of the matrix **M** given by (9b). Throughout we assume that this process is supercritical, i.e., (10) holds. Then branching-process theory (e.g. Harris, 1963) states that the probabilities *σ*_1_ = 1 − *π*_1_ and *σ*_2_ = 1 − *π*_2_ of loss of type1 and type-2 individuals, respectively, are given by the uniquely determined solution (*σ*_1_, *σ*_2_) = (*s*_1_, *s*_2_) of

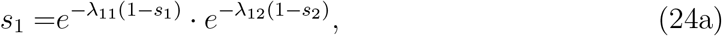

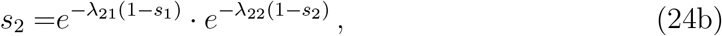

that satisfies 0 < *s*_1_ < 1 and 0 < *s*_2_ < 1. Therefore, the invasion probabilities *π*_1_ and *π*_2_ of *A*_1_*B*_1_ and *A*_1_*B*_2_, respectively, as well as the mean invasion probability 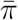 of *A*_1_, defined in equation (7), can be computed numerically. Below, we refer to this as the (exact) numerical solution.

### Remark 7.

i. If *m* = 0, then (9b), (6), and (7) show that *λ*_12_ = 0 and 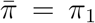. Therefore, by (24a), 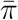 depends only on *λ*_11_, which is independent of *r* if *m* = 0. Thus, 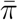 is independent of *r* if *m* = 0. The simple intuitive explanation is that in this case allele *B*_1_ is fixed on the island under our initial equilibrium condition (see eq. 6) so that the mutant *A*_1_ occurs always on the background *B*_1_, which makes recombination irrelevant in the early phase of invasion.
ii. If *r* = 0, then *λ*_12_ = *λ*_21_ = 0 and the principal eigenvalue of **M** becomes *λ* = *λ*_11_ = 1+(*a*+*e*)/(1+*b*). Therefore, *π*_1_ depends only on *λ*_11_, and *π*_2_ depends only on *λ*_22_ = (1 + *a*)/(1 + *b*), which is less than 1 because we assume *a < b*. Therefore, *π*_2_ = 0 and 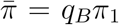, which depends only on *a* + *e* but not on *a* and *e* separately. Also this result has a simple intuitive explanation: In the absence of recombination, the mutant cannot invade if it occurs on the ‘bad’ background *B*_2_, because on this background its invasion rate is *λ*_22_ < 1 and without recombination it cannot recombine onto the ‘good’ background *B*_1_. Because the invasion rate of *A*_1_*B*_1_ is *λ*_11_ = 1 + (*a* + *e*)/(1 + *b*), the migration rate enters the invasion probability 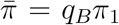 only through the equilibrium frequency of *B*_1_.

In general, the transcendental equations in (24) cannot be solved analytically (see Appendix E for explicit solutions in two cases). However, we can obtain approximations of the invasion probabilities by assuming a slightly supercritical branching process. For this purpose we assume that the mean matrix **M** = **M**(*ϵ*) depends on a parameter *ϵ* > 0 such that its leading eigenvalue, given in (11), can be written as *λ*(*ϵ*) = 1 + *ρ*(*ϵ*), where *ρ*(*ϵ*) → 0 as *ϵ* → 0.

Let *u*(*ϵ*) = (*u*_1_(*ϵ*), *u*_2_(*ϵ*)) and *v*(*ϵ*) = (*v*_1_(*ϵ*), *v*_2_(*ϵ*))^*T*^ be the positive left and right eigenvectors of **M**(*ϵ*) corresponding to the leading eigenvalue *λ*(*ϵ*). They are normalized such that

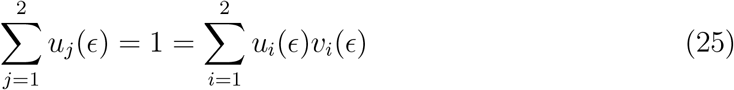

holds. For a slightly supercritical branching process, the invasion probabilities *π*_*i*_ are of order *ϵ* and can be expressed as *π*_*i*_ = *π*_*i*_(*ϵ*) + *o*(*ϵ*), where

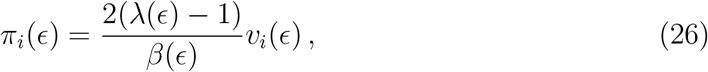

and *β*(*ϵ*) is a complicated expression (Athreya, 1993); see Haccou et al. (2005, pp. 126-128) for a simpler presentation. This approximation is valid for general offspring distributions, such that the matrix **M**(*ϵ*) is primitive for every *ϵ* > 0, i.e., some power is strictly positive. Under the assumption of independent Poisson offspring distributions, we obtain (Appendix B)

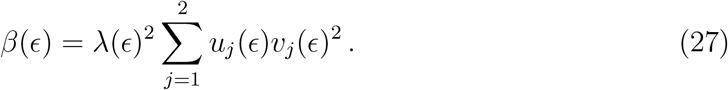

We denote the resulting slightly supercritical approximation for the true mean invasion probability 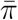 by

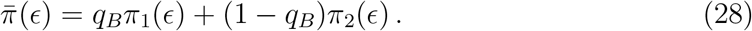

Thus, the argument (*ϵ*) always signifies the approximation.

### Remark 8.

We refrain from giving explicit expressions of quantities such as **M**(*ϵ*), *β*(*ϵ*), or 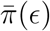 because they are very complicated. They could be obtained, for instance, by setting *m* = *m*\* − *ϵ* since we have noted above that *ϵ* = 0 if *m* = *m*\*. In many of our numerical examples we have *m*\* < 0.1, which would imply *ϵ* < 0.1 because *m* > 0. In addition, *λ* is strictly declining as a function of *r* (File S1, Sect. 6) which assumes its maximum 1 + (*a* + *e*)/(1 + *b*) at *r* = 0 (Remark 7(ii)) and decays to the value 1 as *r* increases to *r*\* (see above). Therefore, the condition of supercriticality will be satisfied if (*a* + *e*)/(1 + *b*) = *O*(*ϵ*) or, simpler, if *a* = *O*(*ϵ*) and *e* = *O*(*ϵ*). Again, rewriting the relevant expressions in terms of *E* makes them very cumbersome. Therefore, in the sequel, we indicate the dependence of *E* only to signify that we use the assumption of slight supercriticality.

### Remark 9.

Aeschbacher and Bürger (2014, eq. (64) in File S1 of their Supporting Information) provided a formula equivalent to

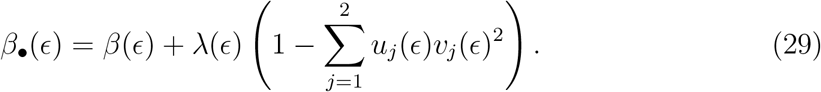

However, in general, *β*_•_(*ϵ*) differs from *β*(*ϵ*). The reason is a trivial calculation error (a lost square) in their derivation of *β*_•_(*ϵ*) from eq. (5.81) in Haccou et al. (2005). Nevertheless, *β*_•_(*ϵ*) yields an approximation 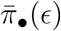 for the true mean invasion probability that has its merits (see below).

### 4.1 Properties of the mean invasion probability 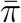 and its approximations

Now we apply the above approximation to our mean matrix (9b) to investigate the properties of 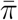 in dependence on the parameters *a*, *b*, *e*, *r*, and *m*. The resulting explicit expressions for the leading terms *π*_*i*_(*ϵ*) and 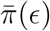, obtained by substituting the explicit expression *β*(*ϵ*) from (27) into (26), are complicated and not very informative (see, e.g., (11) for the principal eigenvalue *λ*). They are derived and presented in File S3, Sect. 2. For the following two special cases, we obtain very simple explicit approximations.

In the absence of migration (*m* = 0), we obtain the approximation

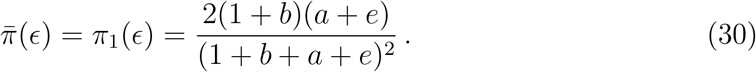

Under the assumption of very weak selection on *A*_1_ (*a*+*e* ≪ 1) this simplifies to 2(*a*+*e*), in analogy to Haldane’s classical approximation. In the absence of recombination (*r* = 0), we obtain

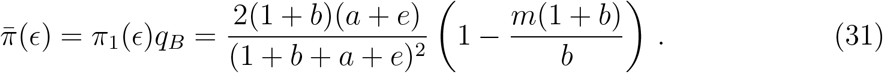

Under the assumption of very weak selection and migration, this simplifies to 2(*a* + *e*)(1 − *m/b*).

These approximations are readily derived from Remark 7(i)and (ii), respectively, and the corresponding first-order approximations of *π*_1_(*ϵ*), eqs. (C.1) and (C.3), in Appendix C and File S3, Sects. 2, 6). An exact expression for the invasion probability 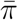 if *r* = 0 is given in (E.2) in Appendix E. It involves the product logarithm. Multiplying this by *q*_*B*_ yields an exact expression for 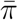 if *m* = 0. Series expansions of these quantities are given in (E.4) and (E.5).

The first-order approximation 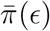 near *m* = 0 is given by eq. (C.4a) in Appendix C and is decreasing in *m*. We conjecture that 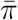 is generally decreasing in *m*, because this is supported by all our numerical results. However, as is clearly visible in Fig. 2 and shown analytically below, 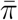 and its approximations are not necessarily decreasing in *r*. This shows that the monotonicity properties of *λ*, which is strictly decreasing in *m* and in *r* by Remark 3(iii), do not necessarily carry over to 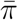.

**Figure 1:**
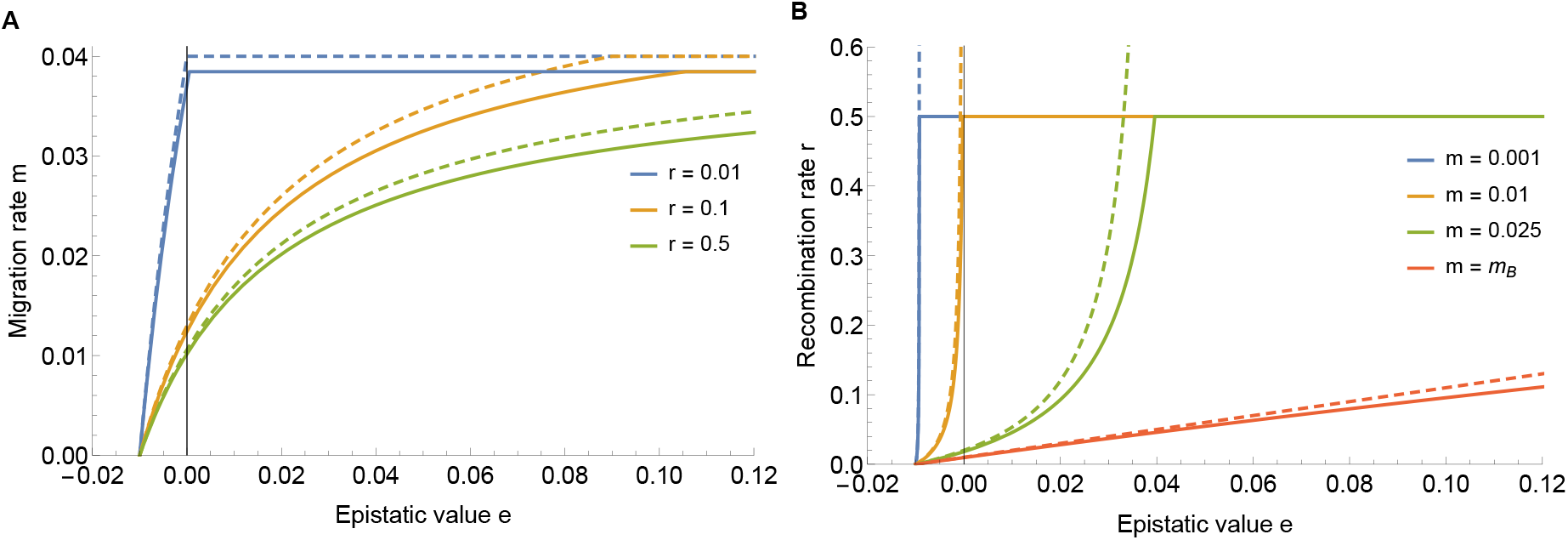
Upper bounds for invasion of *A*_1_ as functions of the epistatic value *e*. In panel A, the upper bound is given by max{*m*\*, *m*_*B*_}, where *m*_*B*_ ≈ 0.0385 because *b* = 0.04; in panel B, it is given by 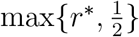. In both panels, the upper bounds are shown as solid curves. Invasion is impossible if *e < −a* = −0.01 and above the solid curves. In panel A and if *r* = 0.5 (green curve), *m*\* < *m*_*B*_ for the whole range of *e*. If *r* = 0.1 (orange) or *r* = 0.01 (blue), then *m*\* < *m*_*B*_ only if *e < e*_*r*_, where *e*_*r*_ ≈ 0.106 or *e*_*r*_ ≈ 0.0005, respectively. In panel B, the critical values *e*_*m*_ below which invasion is restricted to 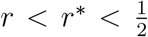 are *e*_*m*_ = −0.0093, −0.002, 0.0396, 1.03 for *m* = 0.001, 0.01, 0.025, 0.0385, respectively. If *e> e*_*m*_, then invasion can occur for every recombination rate. The dashed curves show the corresponding weak-forces approximations, given by (23b) and (23d).

**Figure 2:**
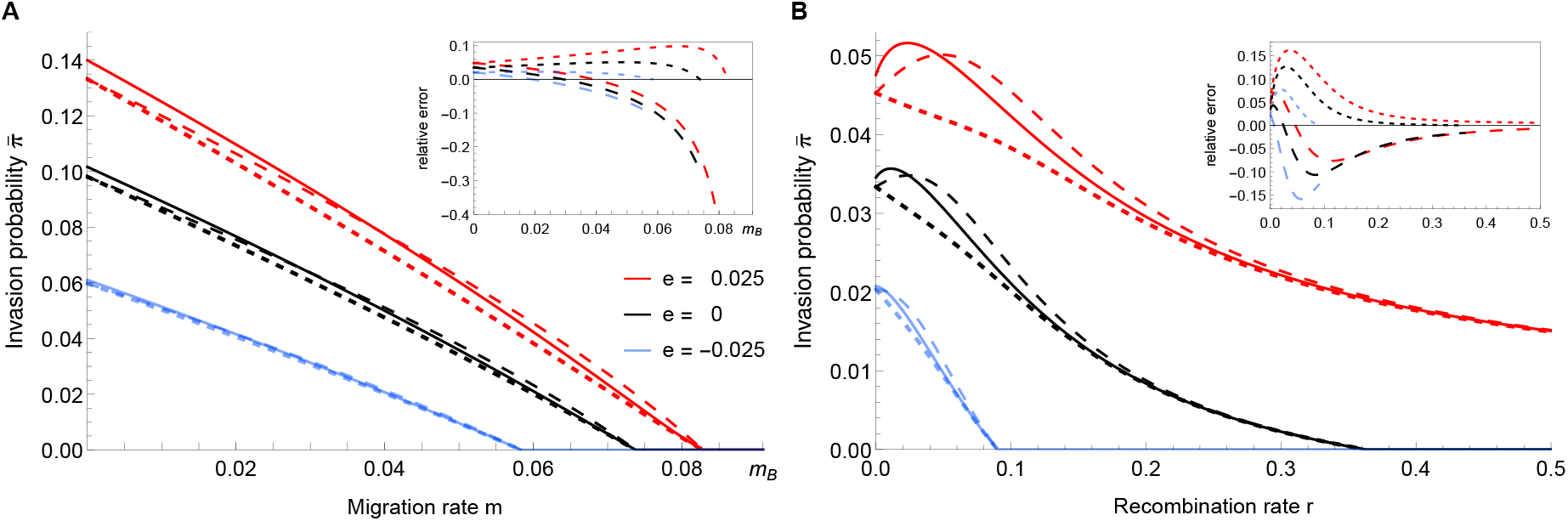
Dependence of the mean invasion probability 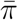 on the migration rate (A) and the recombination rate (B) for three different values of epistasis. The solid curves show 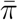, where the numerical solution (*π*_1_, *π*_2_) of the exact equations (24) has been substituted into (7). The dotted curves show the approximation 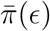, and the dashed curves show 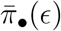 (File S3.2). The line color legend in A applies to both panels. The insets show the relative error of the approximations with respect to the numerical solution of (24), i.e., 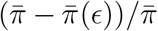. Relative errors are shown only for *m < m*\* (in A) and for *r < r*\* (in B) because 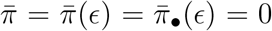 above these critical values, so that the relative error is undefined. The dashing of the curves in the inset corresponds to the dashing of the approximations in the main figure. In both panels we have *a* = 0.06 and *b* = 0.1. In panel A, *r* = 0.1 and in panel B, *m* = 0.06.

Figure 2 demonstrates that both approximations, 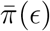 and 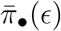, are quite accurate. In fact, in can be shown that they are identical if *m* = 0, *m* = *m*\*, *r* = 0, and *r* = *r*\* (Appendix B). Whereas 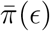 appears to be always smaller than the exact value of 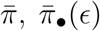 can be smaller or larger. The parameters used in Fig. 2A imply *e*_*r*_ ≈ 0.06. Because *e < e*_*r*_ for all three choices of *e*, all curves decay to 0 in the interval (0, *m*_*B*_) by Proposition 1. The critical values above which invasion becomes impossible are *m*\* ≈ 0.083 for the red curves, *m*\* ≈ 0.074 for the black curves, and *m*\* ≈ 0.058 for the blue curves. In Fig. 2B we have *e*_*m*_ ≈ 0.004. From Proposition 2 we know that epistasis is sufficiently strong to enable invasion for every recombination probability (including free recombination) if and only if *e > e*_*m*_. This is the case for the red curves, where *e* = 0.025. In the absence of epistasis (black curves), we have 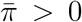 only if *r < r*\* ≈ 0.36, and for *e* = −0.025 (blue curves), we have 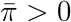 only if *r < r*\* ≈ 0.09.

Figure 2 shows that increasing epistasis (larger *e*) increases the mean invasion probability of *A*_1_, and all our numerical results confirm this. A general proof seems difficult, but at least the principal eigenvalue *λ* is increasing in *e* by Remark 3. With the scaling of epistasis as in (1), this is not surprising because increasing *e* raises the fitness of *A*_1_*B*_1_. Whereas for tightly linked mutants *A*_1_ positive epistasis has only a quantitative effect by increasing the invasion probability, for loosely linked or unlinked mutants (*r* close to 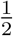) it may have a qualitative effect by enabling invasion.

The weak-forces approximations of the invasion probabilities are simpler than those used above, but still quite complicated. They are provided in Appendix D.

For weak migration and if *A*_1_ alleles have a small effect and are loosely linked to the existing polymorphism, a simple and useful approximation for the invasion probability is obtained. Indeed, if we assume *a, m, e* ≪ *r, b* and use either 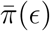 or 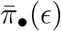, we find to leading order in *a*, *m*, and *e*:

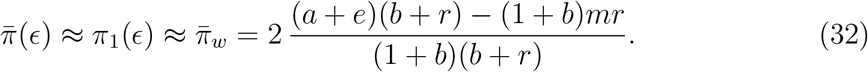

The reason why 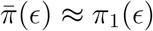 is that if *a, m* ≪ *b*, the frequency of *B*_1_ at the equilibrium *E*_*B*_ is close to one. In addition, the invasion probability *π*_2_ of *A*_1_ on the deleterious background *B*_2_ is always smaller than *π*_1_. Therefore, the contribution of the term (1 − *q*_*B*_)*π*_2_(*ϵ*) in (7) to 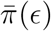 is negligible in this case.

The approximation 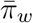 in (32) is increasing in *a* and *e*, and decreasing in *m* and *r*. It shares these properties with the invasion condition (Remark 3) because 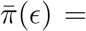 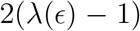 to first order in *a, m, e* if *a, m, e* ≪ *r, b*. If *e* = 0, the approximation (32) simplifies to equation (4) in Yeaman et al. (2016).

From (32), we immediately derive the following approximation for the effect of epistasis on the invasion probability:

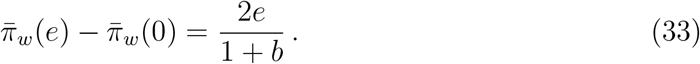

Here, 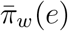 indicates the dependence on *e*. In fact, 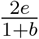 seems to be an upper bound for the increase of the invasion probability caused by epistasis; it is most accurate for very small *m* (Fig. 3).

**Figure 3:**
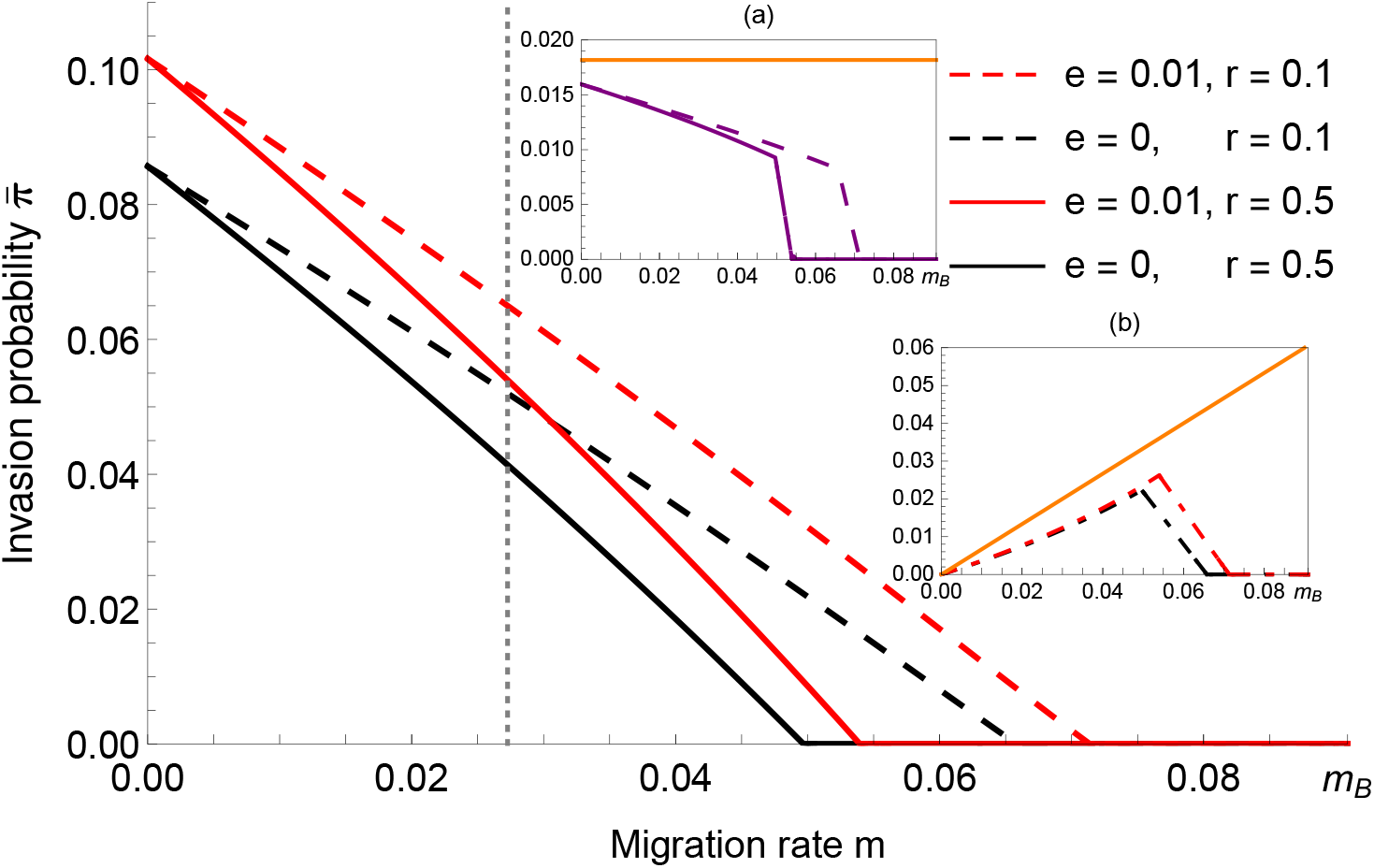
Comparison of the effects of epistasis with those of linkage on the mean invasion probability 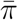, which is shown for the four possible combinations of the two values of *e* and the two values of *r* shown in the legend. The parameters *a* = 0.05 and *b* = 0.1 are kept constant. The figure not only confirms that positive epistasis and linkage increase 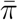 relative to no epistasis and no linkage, but in particular shows that the red solid and the black dashed curves intersect. Therefore, epistasis may be more efficient than linkage in increasing 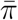 if migration is weak, whereas tighter linkage may be more efficient if migration is strong. The vertical dotted line shows the analytical prediction, 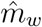 in (37), for this intersection point. Curves of the same color, i.e., with the same *e*, coincide at *m* = 0 by Remark 7(i) and assume the approximate values (30). Inset (a) shows the difference between the two curves of the same line style (dashed or solid), i.e., the effect of epistasis on 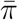. Inset (b) shows the difference between the two curves of the same line color, i.e., the effect of linkage on 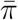. The orange curve in each inset represents the analytical prediction of the respective difference as given in (33) and (34), respectively.

Similarly, the effect of linked relative to unlinked loci can be estimated by

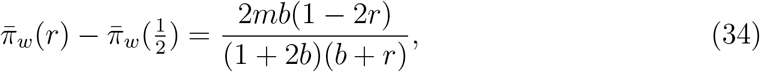

whereas

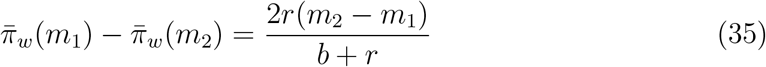

yields the effect of differences in the migration rate. The expressions (33), (34), and (35) give crude but simple approximations for the effects of varying a single parameter (see the orange curves in the insets of Figs. 3 and 4).

**Figure 4:**
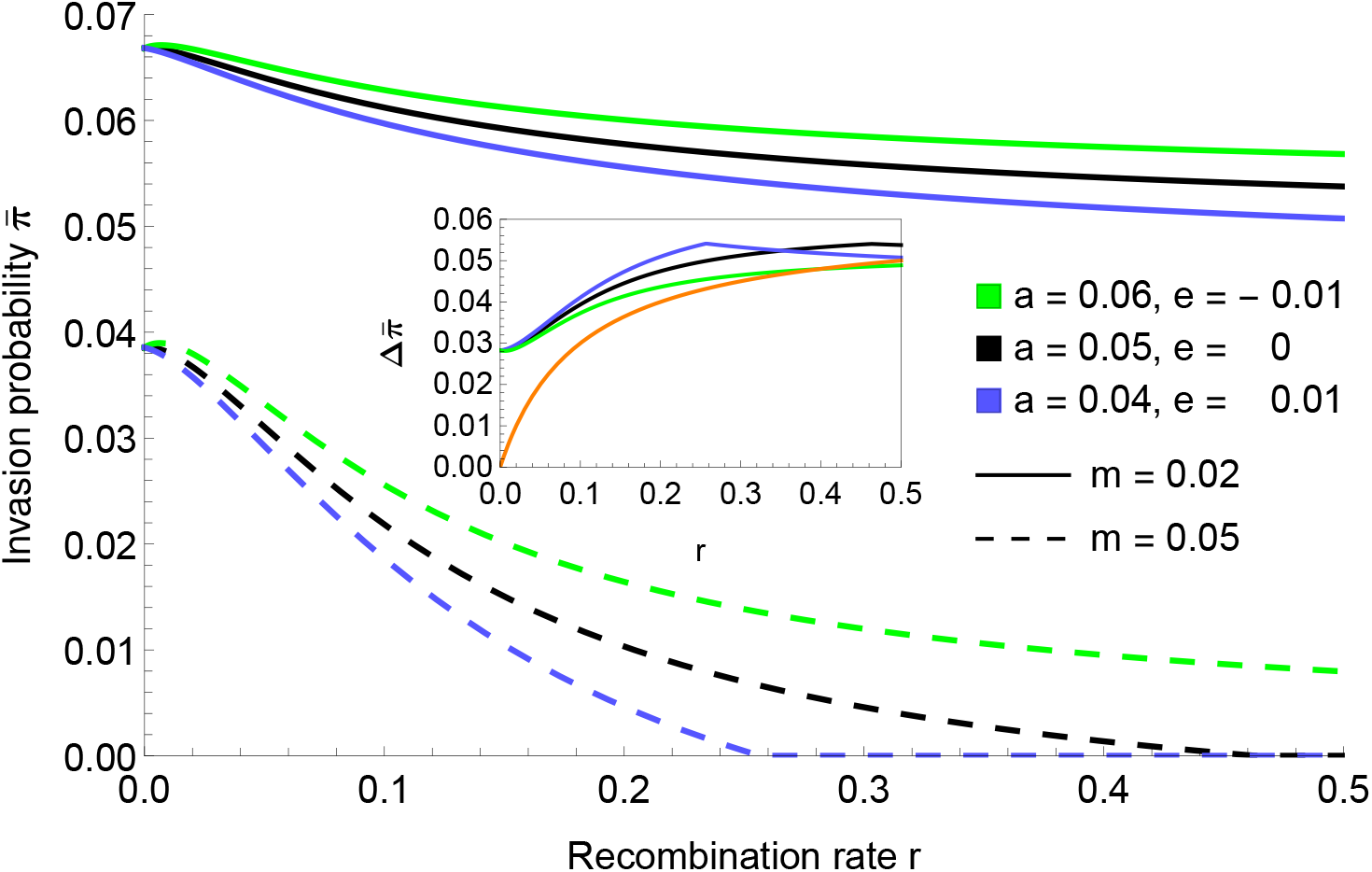
The mean invasion probability as a function of the recombination rate *r* under an alternative parameterization of epistasis. Here, *b* = 0.1 is constant as in the other figures, but *a* and *e* are varied such that the fitness of the best genotype, *A*_1_*B*_1_, is held constant at 1 + *a*′ + *b* = 1 + *a* + *e* + *b* = 1.15. Hence, the fitness of *A*_1_*B*_2_ varies. The inset shows the difference between curves of the same color, i.e., the effect of migration if everything else stays constant. The orange curve represents the analytical approximation (35). All other curves show the exact numerical solution obtained from (24).

We have already seen in Fig. 2 that increasing epistasis and decreasing recombination tend to increase the mean invasion probability. Figure 3 also shows this effect, but in addition it demonstrates the following remarkable effect. If migration is weak, then positive epistasis between unlinked loci is more efficient than stronger linkage between non-epistatic loci in elevating the mean invasion probability, whereas if migration is strong, linkage is more efficient (because the red solid curve and the black dashed curve intersect). A reasonably accurate estimate of the critical migration rate at which these curves intersect is obtained by solving the equation

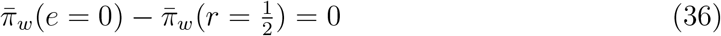

in terms of *m*, which yields

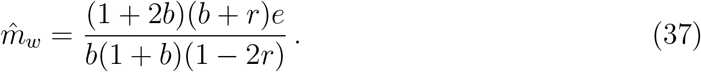

The value of 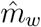 is shown by a vertical dashed line in Fig. 3.

In addition to the estimate (37) of this intersection point, we can derive an exact condition when intersection of two curves, *π*(*a, b, m, e*_1_, *r*_1_) and *π*(*a, b, m, e*_2_, *r*_2_) as functions of *m*, occurs. From either the biological scenario or from inspection of expressions (30) and (E.5), we infer that the invasion probability at *m* = 0 is independent of *r* but not of *e*. Thus, given *a* and *b*, the order of the invasion probabilities at *m* = 0 is solely determined by *e*. If *e* is sufficiently small (*e < e*_*r*_), then *m*\* in (17b) is the upper bound for possible invasion (Proposition 1). Analysis of *m*\* yields the order of the invasion probabilities for large values of *m*. In supplementary file S3 we derive the following condition for an intersection of curves with (*e*_1_, *r*_1_) and (*e*_2_, *r*_2_) in the interval 0 ≤ *m* ≤ *m*\*(*e*_2_, *r*_2_). If

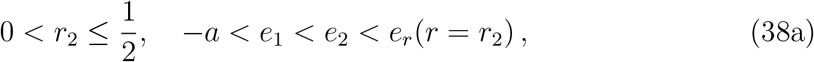

and

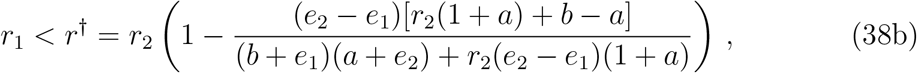

then they intersect. Because *r*^†^ < *r*_2_, the curve corresponding to tighter linked loci and weaker epistasis is indeed above the other if *m* is sufficiently large (but still less than *m*\*). In Fig. 3, we have (*e*_1_, *r*_1_) = (0, 0.1) and (*e*_2_, *r*_2_) = (0.01, 0.5) for the black dashed and red solid curves, respectively. With *e*_1_ = 0, *e*_2_ = 0.01, and *r*_2_ = 0.5, we obtain *r*^†^ ≈ 0.24 *> r*_1_ = 0.1, which explains the intersection.

In Fig. 4, the mean invasion probability 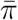 is displayed as a function of *r* for different combinations of *a*, *e*, and *m*. In particular, *a* and *e* are individually varied such that *a* + *e* = constant. Therefore, the fitness of *A*_1_*B*_1_ is fixed for all shown combinations, and only the fitness of haplotype *A*_1_*B*_2_ varies with *a*. The reasoning behind this choice is that, instead of (1), the epistasis parameter *e* could be defined by

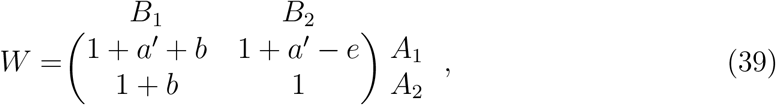

where *a*′ = *a* + *e*. Which scaling is more appropriate may depend on the situation that one intends to model and is discussed below. In light of this parametrization, Remark 7(ii) can be reformulated such that the value of 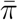 at *r* = 0 is determined only by the parameter *a*′, and not by *e*. For *r* > 0, 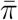 depends on *a*′ and on *e*, as is shown by the divergence among curves of the same dashing style. This is also the case for the first-order approximation of 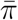 (eq. (C.2) in Appendix C).

Figure 4 shows that the rate of decline of 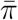 with increasing *r* and constant *a*′ is determined by the value of *e*. A small positive, or even negative value, of *e* entails a slower decay of 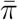, because increasing *e* reduces the fitness 1 + *a*′ − *e* of the haplotype *A*_1_*B*_2_.

The difference of the invasion probabilities for different migration rates (difference between curves of the same color) is shown in the inset. The analytical approximation (35) (orange curve) is quite accurate for strong recombination, but fails for low recombination. This is not unexpected because the approximation (32), from which (35) is derived, assumes large *r*.

Finally, it seems worth noting that 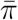 is more sensitive to changes in the recombination rate *r* if *m* is relatively large compared to *a*, as recombination works in conjunction with the afflux of migrant genotypes. In particular, for given selection parameters, the decay of 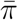 as a function of *r* is faster if *m* is larger (compare the dashed with the solid curves in Fig. 4).

### 4.2 Non-zero optimal recombination rate

The red and black solid curves in Figure 2B demonstrate that the (true) mean invasion probability 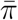 can increase for small *r* and be maximized at a non-zero recombination rate, to which we refer as the optimal recombination rate. First, we derive a (sharp) condition for the existence of a non-zero optimal recombination rate, then we will briefly discuss the approximations shown in the figure.

In addition to our general assumptions (5) and 0 < *a < b*, we assume *a* + *e* > 0 so that the branching process is supercritical for sufficiently small *m* and *r* (Propositions 1 and 2), but not necessarily slightly supercritical. In Appendix E we show that the derivative with respect to *r* of the mean invasion probability 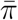 evaluated at *r* = 0 is

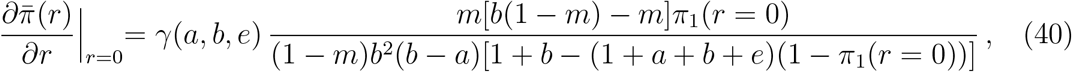

where

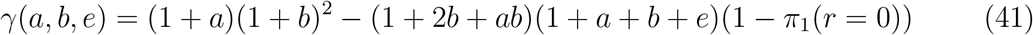

has the same sign as 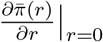 because all other factors are positive. Here, *π*_1_(*r* = 0) is the invasion probability of *A*_1_ on background *B*_1_ in the absence of recombination. Because in this case *λ*_12_ = 0, (24) informs us that *π*_1_(*r* = 0) is the uniquely determined solution in (0, 1) of 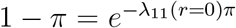, where *λ*_11_(*r* = 0) = 1 + (*a* + *e*)/(1 + *b*). It can be expressed in terms of the product logarithm and has the series expansion

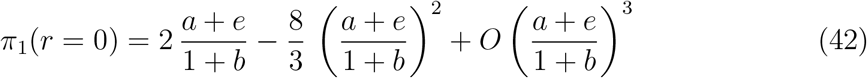

(Appendix E).

Because the function *γ*(*b, a, e*) is strictly monotone increasing in *a*, there exists a unique value *a*\* > 0 for which *γ*(*b, a*\*, *e*) = 0 and *γ*(*a, b, e*) > 0 if and only if *a> a*\*. This critical value *a*\* has the approximation 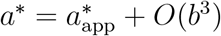, where

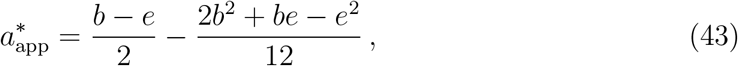

and −*a < e* ≤ *b* is assumed (Appendix E). Thus, increasingly positive epistasis favors the existence of a positive optimal recombination rate because *a> a*_*_ has to be satisfied.

For the parameters in Fig. 2B, where *b* = 0.1 and *a* = 0.06, the critical values are as follows.

- If *e* = 0.025 (red curves), then *a*\* ≈ 0.03582 and 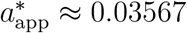,
- If *e* = 0 (black curves), then *a*\* ≈ 0.04844 and 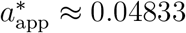,
- If *e* = −0.025 (blue curves), then *a*\* ≈ 0.06118, 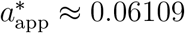.

In accordance with these values, the red and black solid curves in Fig. 2B exhibit a non-zero optimal recombination rate, whereas the blue curve does not. Interestingly, the Aeschbacher-Bürger approximation 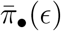 also exhibits a non-zero optimal recombination rate (long-dashed curves in Fig. 2), and yields

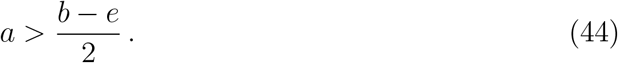

as a condition for it (under the assumption of weak forces; File S5, Sect. 3). Remarkably, this coincides to leading order with (43).

The condition (44) can be rewritten as *b* − *a < a* + *e*. In terms of the fitness matrix (1) this is equivalent to *w*_11_ − *w*_21_ *> w*_21_ − *w*_12_. This means that the fitness benefit of having *A*_1_ instead of *A*_2_ on the *B*_1_ background is greater than that of having the genotype *A*_2_*B*_1_ instead of *A*_1_*B*_2_. If *A*_1_ appears initially on the *B*_2_ background but then recombines onto the *B*_1_ background, this will entail an increase in the frequency of *A*_1_*B*_2_ at the cost of *A*_2_*B*_1_. This is in line with an intuitive explanation for the condition *a> a*\* given in Aeschbacher and Bürger (2014). If *a > a*\*, then *A*_1_ may survive long enough on the bad background *B*_2_ to be rescued to the good background *B*_1_ by recombination. In this case, *π*_2_ (which increases in *r*) provides an important contribution to 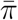 and outweighs the decrease of *π*_1_ as *r* increases.

## 5 Averaged invasion probabilities

Efficient local adaptation of the island population will require the successful invasion, or establishment, of several, if not many, weakly beneficial mutants. They can occur at an arbitrary physical distance to the already established polymorphism, and neither their additive nor their epistatic fitness effects will be known a priori (and often also not a posteriori). Therefore, we investigate various scenarios in which the recombination rate between loci *A* and *B* and the effects *a* and *e* of the new mutants are drawn from appropriate distributions. To obtain expected invasion probabilities of the new mutant *A*_1_, we need to integrate the mean invasion probability 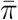 with respect to the chosen distributions of the parameters *r*, *a*, and *e*.

For the recombination rate between *A* and *B* we follow Yeaman et al. (2016) and assume a uniform distribution of 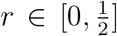; we denote the corresponding probability density by *u*(*r*). For the distribution of (beneficial) additive fitness effects *a*, we assume an exponential distribution (Tataru et al., 2017) with mean 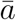, which we denote by

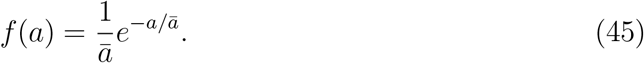

For the distribution of epistatic effects, we consider two cases. In the first, we follow Martin et al. (2007), who deduced an epistasis distribution from an extended version of Fisher’s geometric model and fitted it to two empirical data sets from haploid organisms. Their epistasis distribution is a normal distribution with mean 0 and variance twice the variance of the additive fitness effect of the new mutation (thus, independent of the ‘ancestral strain’), i.e.,

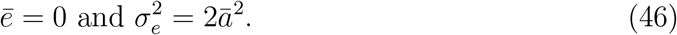

We write *h* for the corresponding probability density function. Defined in this way, *h*(*e*) is a distribution of epistatic interactions between random mutations (i.e., beneficial or deleterious), whereas in our model both mutations (*A*_2_ → *A*_1_ and *B*_2_ → *B*_1_) are beneficial.

Building on the work by Martin et al. (2007), Blanquart et al. (2014) provided results for the mean and variance of a distribution of genetic interactions between beneficial mutations *after* selection. By assuming Fisher’s geometric model in *n* dimensions and a wild type sufficiently far away from the optimum, they showed that the distribution of these (already selected) beneficial mutations is approximately normal with mean and variance given by

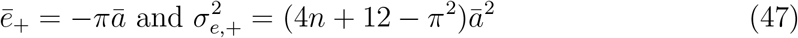

(their eq. 4 applied to our mutation distribution). In the absence of any information on the distribution of epistatic effects of beneficial mutations before selection, we use this distribution in our numerical investigation, mostly for the case *n* = 1. Then 10.2% of all mutations have a positive epistatic effect and 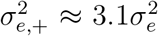. We denote the normal density corresponding to (47) by *h*_+_.

Another caveat is that our model is not based on Fisher’s geometric model. Martin et al. (2007) and Blanquart et al. (2014) assume that the wild type is sufficiently far away from the optimum, such that mutations have small effects compared to the distance from the optimum. In addition, log-fitness is concave, i.e., there is not only directional selection but also a significant component due to stabilizing selection. Therefore, it is expected that beneficial mutations exhibit mostly negative epistasis. Our model can be embedded into their model if the fitness of the island type (*A*_1_*B*_1_) remains lower than the maximum achievable fitness, i.e., more than just two mutations are needed to get close to the maximum fitness. The unboundedness of our distribution *f* will not cause any problems as long as its mean is small enough, so that large effects that overshoot the hypothetical optimum are extremely rare. Therefore, in the figures demonstrating the consequences of drawing the fitness effects from distributions, we assume 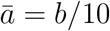. Then additive effects *a* larger than *b* are extremely rare. If directional selection dominates and any stabilizing component can be ignored, then log-fitness will be close to linear and average epistasis will be close to zero.

Because analytical evaluation of the expectations of the approximate invasion probability 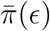 with respect to the distributions of *r*, *a*, or *e* seems unfeasible, we present only results from numerical integration of the exact, numerically determined 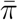. Our main focus is on the role of linkage and epistasis.

We use notation such as

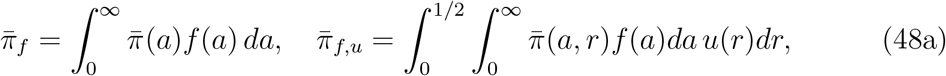

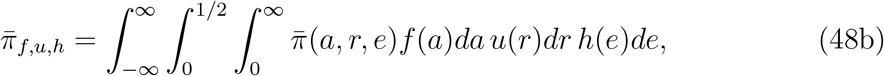

for invasion probabilities averaged over the indicated distributions. To emphasize that a particular value is fixed, e.g., *e* = 0 or 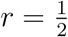, we indicate this by additional subscripts, e.g., by 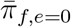 or 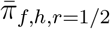, respectively. Since the distributions are independent, a different order of the subscripts does not change the result.

To approximate, for instance, the integral 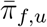, we define a sufficiently fine grid (*a, r*). At each grid point the numerical value of the integrand is computed. The resulting table of values is approximated by a continuous (bivariate) function with the built-in *Mathematica* method Interpolation. This continuous function is then integrated numerically over *a* and *r* (again using *Mathematica*) to obtain an approximation for 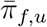.

Figure 5A shows that linkage affects the averaged invasion probability in essentially the same way for all three values *e* = −0.005, 0, 0.005 (compare the shapes of the curves of the same line style but of different color). The averaged invasion probability is always declining in *m*, and for large *r* this decline is much steeper than for small *r*. Indeed, when *m* gets close to *m*_*B*_, invasion probabilities group together that are computed with the same strength of linkage (see inset), whereas for small *m* invasion probabilities based on the same *e* are similar, and identical if *m* = 0 (Remark 7(i)). Because *a* is drawn from the exponential distribution *f*, mutants of large effect can occur and invasion becomes possible for every *m* ≤ *m*_*B*_. However, for large *m* the invasion probability may be negligibly small, especially if *r* is large (see inset). Thus, in concordance with previous results (e.g., Fig. 1B and Fig. 4), this figure confirms that tight linkage becomes essential for facilitating, or even enabling, invasion if migration rates are high.

**Figure 5:**
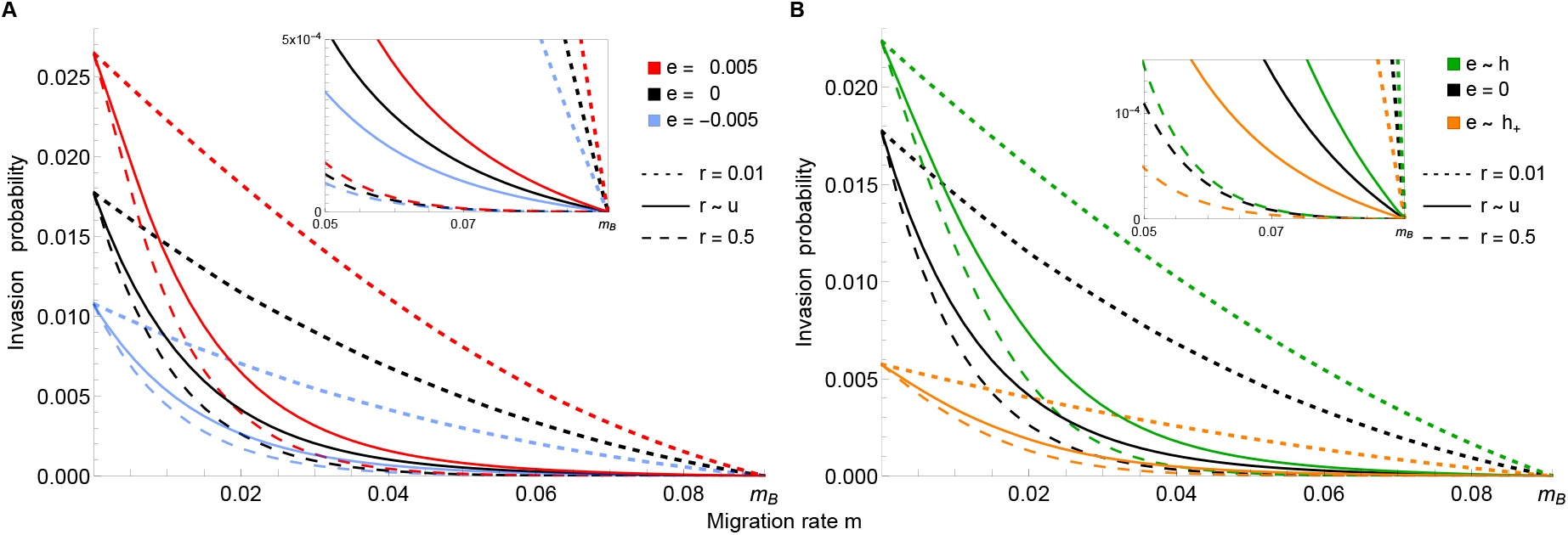
Averaged invasion probabilities of new mutations as functions of *m*. In all cases, the invasion probability is averaged with respect to *f* (*a*) with mean 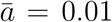. The parameter *b* is fixed at *b* = 0.1, so that the maximum possible migration rate is *m*_*B*_ ≈ 0.091. (A) For each of the (fixed) epistatic values *e* = 0.005, 0, −0.005, the averged invasion probabilities of *A*_1_ under free recombination 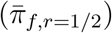 and under tight linkage 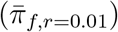 are compared with that under a uniform distribution 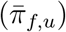, indicated in the legend by *r* ~ *u*. The inset shows the approach to zero for *m* close to *m*_*B*_. (B) is analogous to (A) except that *e* is drawn from the distributions *h* (indicated by *e* ~ *h*, green) or *h*_+_ with *n* = 1 (*e* ~ *h*_+_, orange). Here, 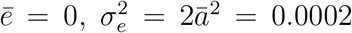, and 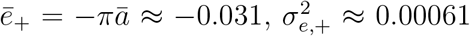 by (46) and (47), respectively. The black curves in A and B (for which *e* = 0) are identical.

Figure 5B demonstrates the consequences of drawing epistatic effects from either the distribution *h* (green curves) or *h*_+_ (orange curves). In the first case, where 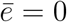, the invasion probability close to *m* = 0 is higher than if *e* = 0, regardless of the degree of linkage (cf. Remark 7). In the second case, in which epistatic effects are negative on average, i.e., 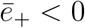, the invasion probability (orange curves) for small migration rates is reduced substantially compared to both other scenarios.

Notably, in Figs. 5A and 5B, curves of different color and dashing style can cross. For instance in B, tight linkage (short-dashed black curve) provides a greater adavantage for invasion than high fitness induced by epistasis (solid and long-dashed green curves), unless *m* is very small. Similarly, above a critical migration rate, the invasion probability of tightly linked mutations with, on average, negative epistasis (short-dashed orange curve) is higher than that of unlinked or loosely linked mutations with no epistasis (long-dashed and solid black curves), and even higher than that with, on average, positive epistasis (long-dashed and solid green curves). We observed this effect already in Fig. 3 for fixed values of *e* and *r*, where we discussed it briefly. In particular, (37) provides a reasonably accurate approximation for the critical value *m* at which the advantages of higher epistasis and tighter linkage balance.

The effects of linkage and epistasis on the invasion probability are further highlighted in Fig. S3, which is based on the data and graphs in Fig. 5, and in Fig. S5. Both demonstrate that the epistasis distribution *h*, which has mean 0, facilitates invasion of *A*_1_ compared to absence of epistasis, whereas the epistasis distribution *h*_+_, which has a negative mean, impedes it. In addition, Fig. S3B shows that if migration is weak, invasion at unlinked loci contributes more to the average invasion probability than invasion at linked loci, whereas the contribution of linked loci matters most if migration is strong.

## 6 Average effects of successfully invading mutants

If the mutational effects, additive or epistatic, are drawn from distributions, mutants with larger effects will have an elevated invasion probability. However, mutants with much larger effects than the mean will also have a substantially reduced probability of occurrence. Here, we explore the average effects of successfully invading mutants numerically; given the complexity of the approximations for 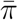 an analytical treatment seems out of reach. In view of the applications in the next section, we study the average effects as functions of the recombination rate between the two loci.

Each beneficial mutant has an additive effect *a*, drawn from the exponential distribution *f* in (45), and an epistatic effect *e* drawn from either the normal distribution *h* in (46) or the normal distribution *h*_+_ in (47). For given parameters *b*, *m*, *r*, and 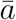 (the mean of *f*) and following the previous notation, we denote the invasion probabilities averaged over *f* and *h* or over *f* and *h*_+_ by

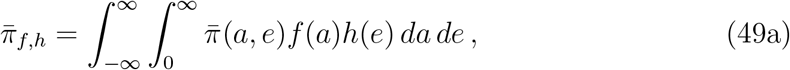

or

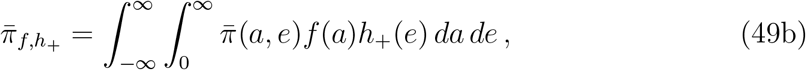

respectively. Then the average effects (*a, e*) of the successfully invading mutations, i.e., conditioned on invasion, are given by

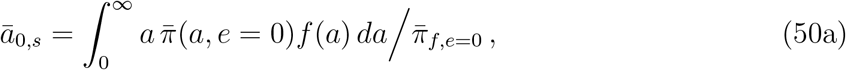

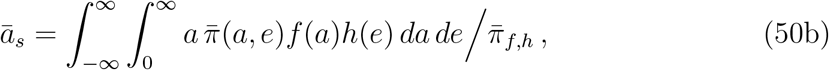

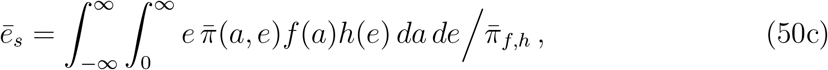

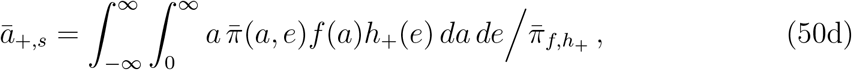

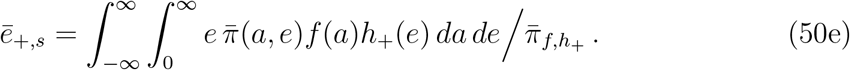

Figure 6 displays these mutational effects for a variety of migration rates as functions of the recombination rate. As is expected intuitively on the basis of the above developed theory, the average additive effects (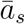 and 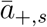) as well as the average total effects (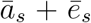 and 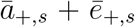), which are expressed in the island genotype *A*_1_*B*_1_, always increase with increasing migration rate and with increasing recombination rate. This is because the minimal effects of *a* and *e* necessary for invasion increase with increasing *m* or *r*, as follows from Remark 3(ii) or 3(iii), respectively. However, the corresponding invasion probabilities (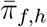 and 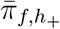) decline (often rapidly) with increasing *m* or *r* (Fig. S5). Thus, the waiting time until a mutation becomes established at a large recombinational distance from the already polymorphic locus may be very long. If it becomes established, then it likely is a mutation of large effect.

**Figure 6:**
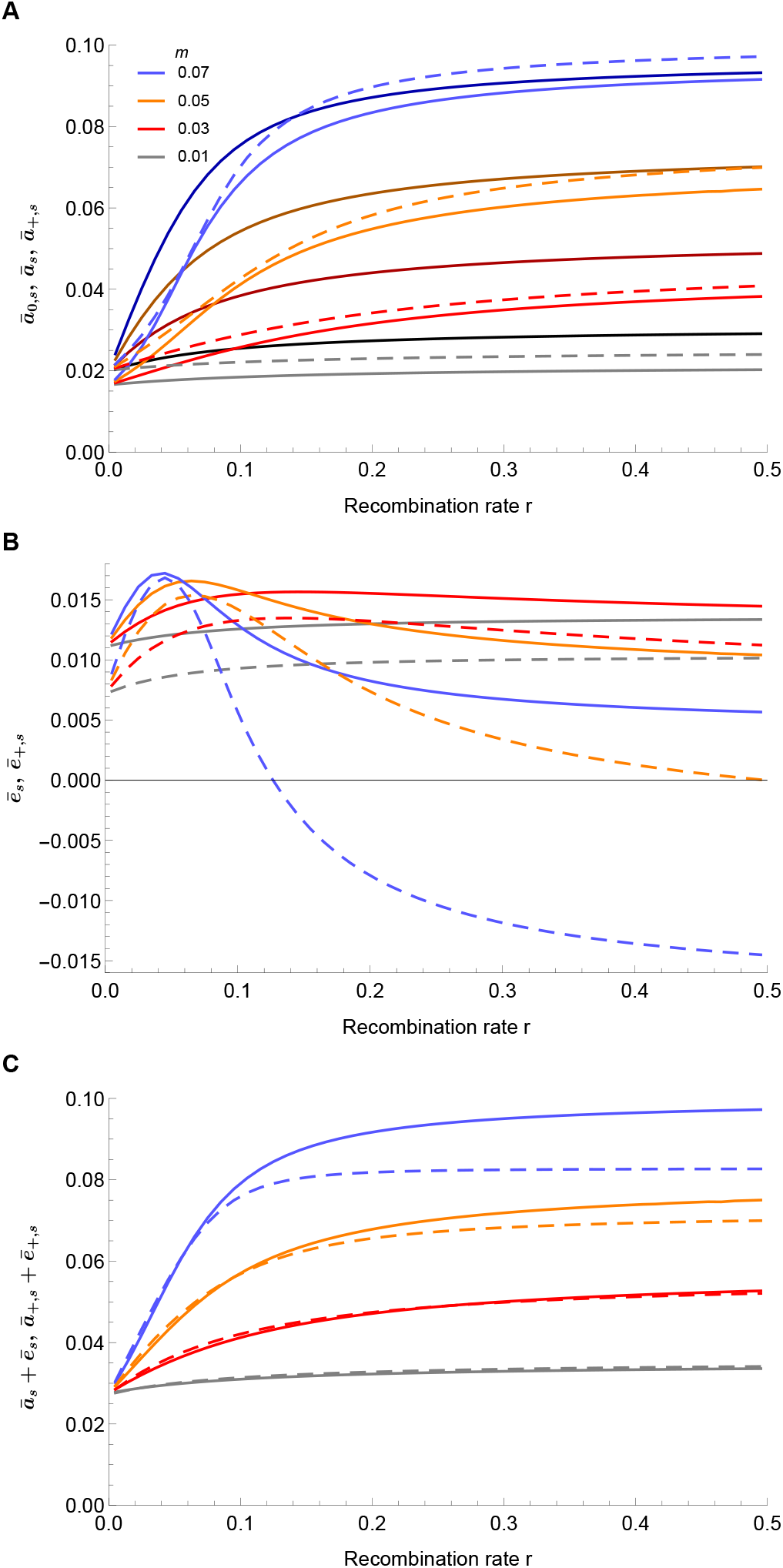
Average additive and epistatic effects of successfully invading mutants as functions of the recombination rate *r*. Additive effects are drawn from an exponential distribution with mean 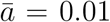, epistatic effects from *h* in (46) or *h*_+_ with *n* = 1 in (47). Panel A shows the resulting average additive effects 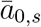 (solid, darker color), 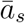 (solid, lighter color) and 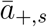 (dashed) for the indicated migration rates. For unlinked, non-epistatic mutations (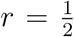, *e* = 0) the minimum effects required for invasion are *a* ≈ 0.0093 (if *m* = 0.01), *a* ≈ 0.0291 (if *m* = 0.03), *a* ≈ 0.0504 (if *m* = 0.05), and *a* ≈ 0.0735 (if *m* = 0.07); the values 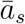 and 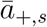 at 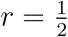 lie well above them. Panel B shows the corresponding average epistatic effects 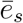 (solid) and 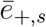 (dashed). Panel C shows the (total) average effects of *A*_1_ in the island genotype *A*_1_*B*_1_, i.e., 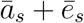 (solid) and 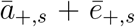 (dashed). In all cases *b* = 0.1. Figure S5 shows the corresponding invasion probabilities.

Panel A shows that 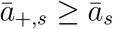 holds always. The reason is that mutants arising under the distribution *h*_+_ have on average negative epistatic effects 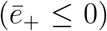, whereas those arising under distribution *h* have 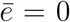. Therefore, as panel B shows, 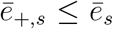 holds and, for the largest migration rate (*m* = 0.07) 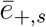 can even be negative. (Recall from (47) that about 10% of the mutants drawn from *h*_+_ have a positive effect.) Because a larger total effect (*a* + *e*) facilitates invasion, 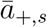 is larger than 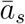. As panel C shows, the average total effects 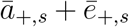 and 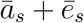 are almost identical for the two smaller migration rates, but differ considerably for the largest migration rate; then 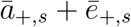 is well below 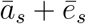 if *r* ≳ 0.12, because then 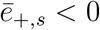 (see panel B).

It is also of some interest to compare these average values with 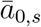, the average values of successful mutations that are not epistatic, i.e., if *a* is drawn from *f* and *e* = 0. For the migration rates *m* ≤ 0.05 in Fig. 6, we observe the following order, valid for every *r*:

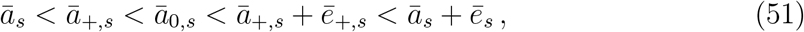

If *m* = 0.7, then 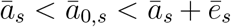 still holds, but the other inequalities are violated if *r* ≳ 0.12.

## 7 Approximate size of a genomic island

Yeaman et al. (2016) discussed several categories of explanation for the occurrence or maintenance of genomic islands of elevated divergence (measured by *F*_*ST*_) between a pair of parapatric populations. A particularly likely explanation is that linkage of locally beneficial de novo mutations to an already established selection-migration polymorphism facilitates successful invasion of such mutations in the face of maladaptive gene flow. Because linkage depends strongly on physical distance, invasions should be successful predominantly locally around the already polymorphic site and thus lead to genomic islands of divergence.

Similar to the quantity *C*_95_, which was introduced by Yeaman et al. (2016), we investigate the quantities *C*_50_ and *C*_90_, the 50% and 90% window sizes. These are the smallest neighborhoods of the polymorphic site in which 50% and 90%, respectively, of all new mutations became established. Thus, *C*_50_ is the value of *r* required to contain 50% of the probability density of 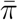 as a function or *r*. Formally, *C*_50_ is defined such that 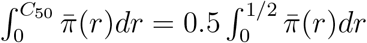 holds, and analogously for *C*_90_.

A small value of *C*_50_ (*C*_90_) corresponds to a locally restricted genomic region in which the mutations profit from their linkage to the polymorphic site. A high value of *C*_50_ (*C*_90_) indicates that loosely linked mutations, i.e., mutations in a large genomic region, contribute substantially to the cumulative invasion probability. Because recombination rates are drawn from a uniform distribution on [0, 0.5], *C*_50_ and *C*_90_ assume the maximum values 0.25 and 0.45, respectively (at *m* = 0).

We compute *C*_50_ and *C*_90_ numerically and focus, in particular, on the effect of epistasis on the window size. In accordance with intuition, *C*_50_ (*C*_90_) decreases with increasing *m* in all observed instances (Figs. 7 and 8). In the absence of epistasis this was also observed for *C*_95_ by Yeaman et al. (2016). The decline in their Fig. 3 looks much sharper than in our Fig. 8, but this is deceptive because they used a logarithmic scale.

**Figure 7:**
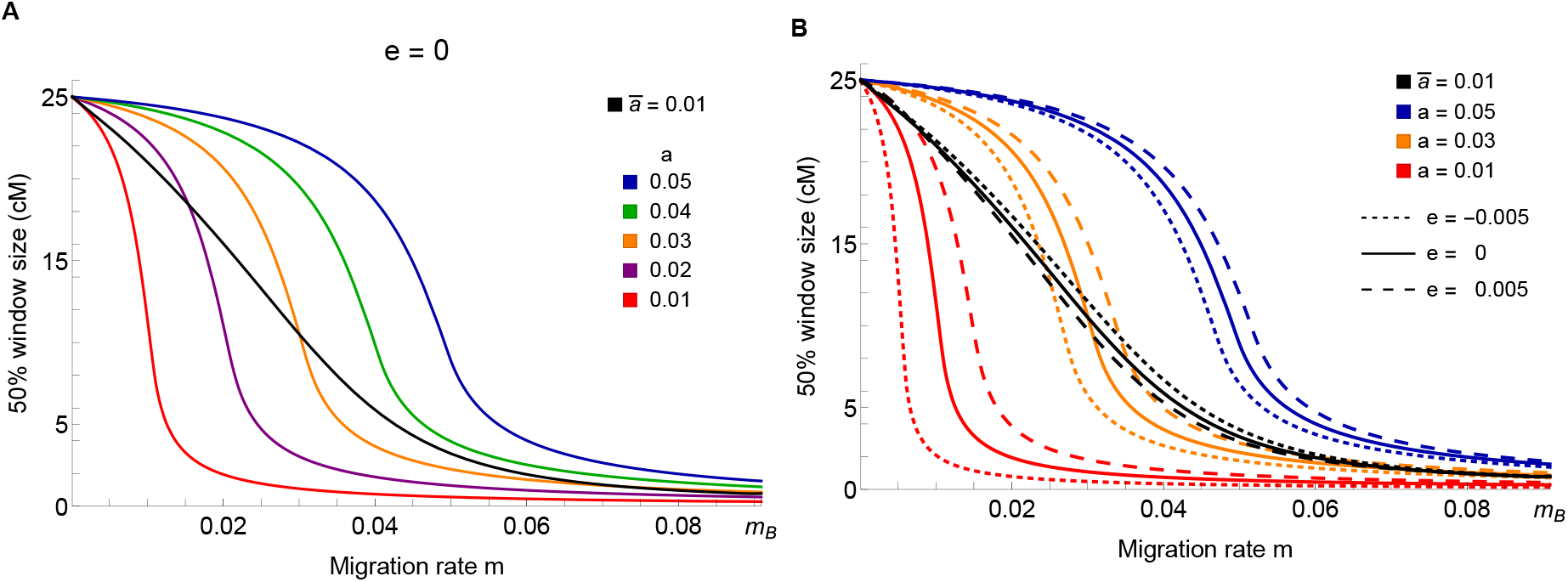
Comparison of *C*_50_ window sizes for five different values of fixed *a* and for *a* drawn from the exponential distribution *f* with mean 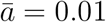. In panel A, absence of epistasis is assumed. In panel B, fixed positive (*e* = −0.005, dashed) and fixed negative epistasis (*e* = −0.005, dotted) epistasis are included and shown for three fixed values of *a*. Note that for *a* drawn from *f*, the curve with negative epistasis (black, dotted) is higher than that without epistasis (black, solid) as well as that with *e* = 0.005 (black, dashed).

**Figure 8:**
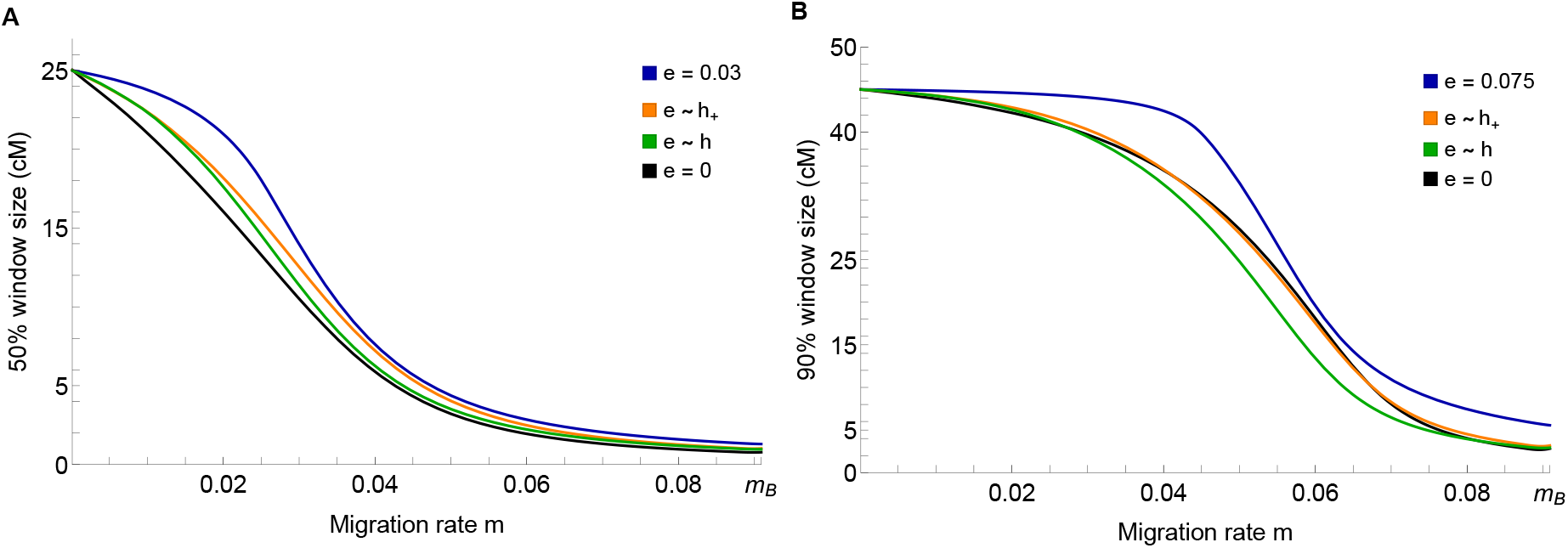
The size of the window within which 50% (A) or 90% (B) of all successfully invading linked de novo mutations occur (*C*_50_, *C*_90_). In all cases *b* = 0.1 and *a* is drawn from an exponential distribution with mean 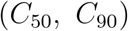. The black curves are for additive mutants and the blue curves are for fixed values of epistasis (*e* = 0.03 in A, *e* = 0.075 in B). The orange and the green curves show the window sizes if the epistatic effects are drawn from *h*_+_ with *n* = 1 and *h*, respectively. The units on the ordinate are in centimorgan, thus we identify *r* = 0.01 with 1 cM. Based on Haldane’s mapping function, this is a suitable approximation if *r* ≲ 0.25.

Figure 7A shows the dependence of the *C*_50_ window size on *m* if epistasis is absent and the additive effects *a* are fixed and increase in a series of small steps (colored curves). These curves differ markedly from the black curve, which displays *C*_50_ when *a* is drawn from an exponential distribution with mean 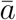. Fixed additive effects lead to a sharp decline in window size in a relatively narrow range of migration rates. This is due to the fact that for increasing *m* the probability of invasion is rapidly decreasing as *r* increases. The much more gradual decline of *C*_50_, when the values of *a* are drawn from *f*, is explained by the fact that with increasing *r* successfully invading mutants have increasingly larger additive effects *a*, as is reflected by the increase of 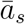 shown in Fig. 6A. Qualitatively, these observations carry over to the cases when *e* is slightly negative or positive and fixed (panel B). The effect of epistasis is strongest in the range of migration rates, where the decline is steepest. Not surprisingly, the effects are stronger for small *a* than for large *a*, because then the ratio |*e*|*/a* is larger. However, the order of the black dashed curves (where *a* is drawn from *f*) is reversed relative to the curves with a fixed value of *a*. If *e* < 0 then the values of *a* that lead to invasion are larger than if *e* = 0 or *e* > 0 (Fig. S6), and larger values of *a* enable invasion in a wider neighborhood of the already polymorphic locus.

Figure 8 demonstrates the effects of choosing the epistatic effects *e* from different distributions, and compares *C*_50_ with the corresponding *C*_90_. Whereas *C*_50_ declines nearly linearly in *m* until *m* ≈ 0.05 and then slowly to near zero at *m* = *m*_*B*_ ≈ 0.09 (Fig. 8A), *C*_90_ declines very slowly between *m* = 0 and about *m* ≈ 0.03 and then rapidly to very small values (Fig. 8B). In each of the two panels, the decline occurs for somewhat larger values *m* if the epistatic value is high and fixed (blue curves), and then declines more sharply than in the other three cases (no epistasis or epistatic coefficients drawn from either *h* or *h*_+_). The reader may keep in mind that the window size is computed based on successfully invading mutants; therefore, the fact that 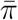 decreases as a function of *m* (Sect. 4) does not directly yield the observed decay of *C*_50_ or *C*_90_.

Somewhat surprisingly, Fig. 8 shows that the window size depends only very weakly on epistasis, unless a fixed high epistatic value is assumed (as for the blue curves). It may seem counterintuitive that *C*_50_ and *C*_90_ are higher if epistatic effects are drawn from *h*_+_, which has the negative mean 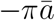, than if they are drawn from *h*, which has mean 0.

These observations have a relatively simple explanation. The distribution *h* has a variance of 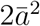, and the variance of *h*_+_ is about 3 times larger (eqs. 46, 47). Therefore, about 10% of mutational effects drawn from *h*_+_ are positive (compared to 50% of those drawn from *h*). This is still enough that for both distributions most of the successfully invading mutants have a positive epistatic effect, often close to 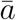 (see Fig. 6B). The fact that the invasion probability of mutants with epistatic effect drawn from *h*_+_ typically is 1/5 - 1/4 of that of mutants with epistatic effect drawn from *h* (see Fig. S5) does not directly affect the window size, which is conditioned on successful invasion. Because successful mutants with *e* drawn from *h*_+_ have a smaller average value than mutants with *e* drawn from *h*, i.e., 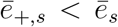, they have a slightly higher additive effect, i.e., 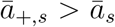 (Fig. 6A). Their total average effects 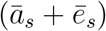 are nearly identical except in the high-migration case *m* = 0.07 (Fig. 6C), where invasion of loosely linked mutations is already extremely unlikely. In summary, mutants with *e* drawn from *h*_+_ can invade at a (slightly) higher distance from the polymorphic site than mutants with *e* drawn from *h*, because invasion of the large-effect mutations under *h*_+_ depends only weakly on linkage. The effects of drawing *e* from a distribution become more pronounced than in Fig. 8 if the variance of *h*_+_ is increased, as is the case if *n* = 3 or *n* = 5 is chosen in eq. (47) (see Fig. S7).

## 8 Discussion

We have performed an analysis of the effects of epistasis on the fate of a new, weakly beneficial mutation in a haploid population that is exposed to maladaptive gene flow. The mutant, *A*_1_, arises at a locus *A* that is linked (with recombination rate *r*) to an already established migration-selection polymorphism *E*_*B*_ at a locus *B*. The existence of the polymorphism requires that the immigration rate *m* of the deleterious allele *B*_2_ is bounded by the constant *m*_*B*_ = *b/*(1 + *b*), where *b* is the selective advantage of *B*_1_. In particular, we characterized the region of the parameter space, in which the de novo mutation *A*_1_ can survive the stochastic phase after its occurrence (Propositions 1, 2, 5, 6). The first two propositions apply to evolutionary forces of arbitrary strength, whereas the two others are derived under the assumption of weak evolutionary forces. This yields simple and intuitive conditions for invasion. In each case, invasion is impossible if *a* + *e* < 0, where *a* > 0 is the mutant’s effect on background *B*_2_ and *a* + *e* that on *B*_1_. In this parameter regime, *A*_1_ exhibits sign epistasis, i.e., the mutation *A*_1_ is deleterious in the presence of *B*_1_, but beneficial on the *B*_2_ background. Therefore, sign epistasis prevents establishment of *A*_1_ independently of the (total) strength of the evolutionary forces. This is in line with the known effect of sign epistasis to constrain the selective availability of mutational trajectories to genotypes of high fitness, as laid out by Weinreich et al. (2005).

If the strength *e* of epistasis exceeds −*a* but is below a certain bound *e*_*r*_ or *e*_*m*_, which depends on *b* and on *r* or *m*, respectively, then invasion is possible below a critical value of *m* or *r*, respectively. If *e> e*_*r*_ or *e > e*_*m*_, then invasion of *A*_1_ in an equilibrium population at *E*_*B*_ is possible for every admissible *m* or *r*, respectively (Propositions 1 or 2). The critical values *e*_*r*_ and *e*_*m*_ can be positive or negative. If they are positive, then an additive mutant *A*_1_ (*e* = 0) can invade only for sufficiently small *m* or *r*, respectively (Fig. 1). In summary, for fixed *b*, increasing *e* (and of course increasing *a*) widens the range of parameters in which invasion can occur.

It is also important to determine if *A*_1_ can enter the population exclusively through the single-locus polymorphism *E*_*B*_, or if there are further possibilities. If allele *B*_1_ is absent from the island population, then, of course, *A*_1_ can invade the resident population consisting of *A*_2_*B*_2_ individuals if 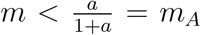. If *B*_1_ is present and 0 < *m < m*_*B*_, then the single locus polymorphism exists and invasion of allele *A*_1_ is possible under the conditions derived in Propositions 1 and 2. If *m > m*_*B*_ > *m*_*A*_ (where the latter inequality holds because we assume *a < b*), then the island population is swamped by the continental type *A*_2_*B*_2_ which eventually becomes fixed.

Intuition about the long-term fate of *A*_1_, and of all genotypes, may be gained from the analysis of the deterministic haploid two-locus two-allele model with continent-to-island migration in Bank et al. (2012). These authors showed that the maximum possible number of internal equilibria is three and at most one is stable. This stable equilibrium corresponds to a so-called Dobzhansky–Muller incompatibility (DMI), which plays a key role in explanations of the evolution of intrinsic postzygotic isolation. In our stochastic model, this DMI will correspond to a quasi-stable state and convergence to it is the most likely outcome after successful invasion. On an extremely long time scale and in the absence of further mutation, fixation of a gamete will occur.

Branching process theory yields a system of equations from which the probability of invasion can be computed. Because they are transcendental, only numerical but no explicit analytical solution is possible. However, by assuming that the branching process is slightly supercritical, analytical approximations become available (Athreya, 1993; Haccou et al., 2005, and Sect. 4). Although the explicit approximations are complex and not directly informative (File S3.2), they are useful to investigate the dependence of the invasion probability on the model parameters (Sect. 4.1). We find that the invasion probability is always increasing with decreasing *m* and with increasing *a* or *e*. For a wide region of parameters, it also decreases with increasing *r*. Nevertheless, there is a significant parameter region in which a non-zero optimal recombination rate occurs, i.e., where the invasion probability is maximized at an *r* > 0 (Sect. 4.2). This phenomenon was already observed in the diploid case without epistasis by Aeschbacher and Bürger (2014). However, with positive epistasis a non-zero optimal recombination rate occurs for a wider range of parameters and the optimum may be even at 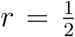 (Figs. 2B and S1).

Related results were found in two-locus models of panmictic populations by Ewens (1967), who used a branching process approximation to study invasion, and by Lessard and Kermany (2012), who used an ancestral selection-recombination graph to study fixation of new mutants. The latter authors showed that in a large (but finite) population, the ultimate fixation probability of a mutant increases if it is in positive epistasis with another beneficial, already segregating mutant, or if it is itself beneficial and there is no epistasis. These results were shown for weak recombination and weak selection and confirm the Hill-Robertson effect. In our branching process model, the Hill-Robertson effect does not seem to be responsible for the non-zero optimal recombination rate, because the population of resident alleles is infinite by assumption.

In our model, low recombination favors the spreading of the mutant if it initially occurs on the good background (*B*_1_), whereas strong recombination is necessary for successful invasion if it occurs on the bad background (*B*_2_). In this case, stronger recombination promotes the escape to the good background. This will lead to successful invasion only if *A*_1_ has a sufficiently high fitness effect *a* on the bad background to survive until it becomes rescued by recombination. Additionally, the effect *a* + *e* on the good background needs to be sufficiently large that invasion can occur despite the higher recombination rate, i.e., *a* + *e > b* − *a* must hold; see (44).

A comparison of the effects of epistasis and linkage on the invasion probability shows that for weak migration a change in epistasis has the stronger effect, whereas for strong migration a change in linkage has the stronger effect (analysis leading to eq. (38) and Fig. 3). In biological terms this can be explained as follows. With weak migration only very few continental *B*_2_ alleles occur on the island. Therefore, recombination occurs mostly between haplotypes with the *B*_1_ allele and has little effect, even if strong. Higher epistasis, however, does always increase the invasion probability. With strong migration, the proportion of recombination events between haplotypes *A*_1_*B*_1_ and *A*_2_*B*_2_ is higher, and the fitness of the recombination products *A*_1_*B*_2_ and *A*_2_*B*_1_ does not depend on epistasis, which reduces the effect of epistasis.

Local adaptation to a new environment is often modeled by assuming that a quantitative trait is subject to Gaussian stabilizing selection, where the population mean is displaced from the fitness optimum. If this displacement is very large, the fitness landscape around the wildtype may close to linear or even convex. In particular, every mutation that brings the population closer to the optimum is beneficial and positive epistasis will accelerate adaptation. For this situation our parametrization in equation (1) is well suited. If the population is not far from the optimum, the fitness landscape will be concave and the maximum fitness cannot be exceeded by new mutations. In this situation, the parameterization in (39) may be most appropriate. If mutations of large effect can overshoot the optimum, then, depending on the background on which they occur, they may reduce fitness. In such a case, negative epistasis may be beneficial for invasion. This effect is demonstrated in Fig. 4, where the highest invasion probability occurs for negative epistasis, especially if linkage is weak (green curves). Once *e* < −*a*, invasion is no longer possible in our model (Proposition 1). Obviously, our model was not designed to study invasion close to a fitness optimum.

Because, in general, neither the fitness effects (*a*, *e*) of mutants nor their recombinational distance (*r*) from the existing polymorphism are known, especially not a priori, in Sect. 5 we studied expected invasion probabilities if *a*, *e*, and *r* are drawn from probability distributions. We assume that the distribution *f* of additive fitness effects is exponential with mean 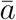. Recombination rates are drawn from a uniform distribution.

The distribution of epistasis (in fitness) depends crucially on the relative position of the population with respect to the optimum on the fitness landscape. We use two different ones. The normal distribution *h*, which has mean 0 (see eq. 46), was derived from Fishers geometric model for the epistatic interaction between arbitrary mutations (Martin et al., 2007). This may be only of limited suitability for our purposes, because we model epistasis as the interaction between *A*_1_ and *B*_1_, which both are advantageous. To account for the interaction of beneficial mutations, we use *h*_+_, a normal distribution with a negative mean (see eq. 47). This distribution was also derived from Fisher’s geometric model by considering the interactions of beneficial mutations (Blanquart et al., 2014). In the derivations of *h* and of *h*_+_, it was assumed that the population is sufficiently far away from the optimum.

We found that a change in the recombination rate affects the invasion probability in a similar way across different epistatic values. Drawing the recombination rate from a uniform distribution again has a similar effect across different epistatic values (Fig. 5). Thus, recombination and epistasis affect the invasion probability essentially independently.

Averaging over epistasis has more subtle consequences for the invasion probability which depend on the properties of the distribution of epistasis. The effects of averaging over *h* or *h*_+_ can be summarized as follows. For each degree of linkage, the larger share of positive epistatic values in the distribution of *h* leads to an increase of the invasion probability compared to the non-epistatic case. The opposite is true if *e* is distributed according to *h*_+_, which has a negative mean. These facts are apparent in Fig. S3B, which shows the ratios of the invasion probability with and without epistasis for the same degree of linkage. Comparison of curves with different degree of linkage in Fig. 5 shows that an effect observed already in Fig. 3 (for fixed values of *e* and *r*) extends to distributions of the parameters *r* and *e*. For small values of *m*, a larger fraction of positive epistatic values is more efficient in boosting the invasion probability compared to a non-epistatic scenario with tighter linkage. This is reversed above an intermediate value of *m* when non-epistatic linked mutations contribute more to the invasion probability (e.g., compare the black solid curve with either the orange dotted or the dashed green curves in Fig. 5B).

Finally, we investigated the effect of epistasis on the size of the genomic window, or neighborhood, around an already existing polymorphism in which 50% or 90% of all successfully invaded linked de-novo mutations occur. For fixed additive and epistatic effects, a sharp decline of the window size occurs in a small range of migration rates, whereas the decline is much more gradual if effects are drawn from a distribution (Figs. 7, 8, S7). If the additive fitness value is drawn from an exponential distribution, then drawing the epistatic value from either of our two distributions does not substantially affect *C*_50_ and *C*_90_ in comparison to *e* = 0. This is in contrast to the distinctively different effects of these distributions on the average invasion probability (Fig. 5B). The reason is that *C*_50_ and *C*_90_ are computed by conditioning on successful invasion. Indeed, the average additive and average total effects of successfully invading mutations are very similar in both cases except for the largest migration rate (Fig. 6), but the respective invasion probabilities may differ by about one order of magnitude (Fig. S5). In summary, unless epistasis is consistently positive and very strong, on average it seems to have a weak effect on the size of genomic islands.

Because sign and strength of epistasis affect the invasion probability of mutants substantially, they will also strongly affect the time horizon in which adaptation occurs and clusters of beneficial mutations, or genomic islands, build up. Analogously, invasion probabilities may decline by several orders as the migration rate increases (Fig. S5), and therefore the emergence of very tightly linked clusters, as predicted by the small window sizes at higher migration rates, will occur only on very long time scales.

The average invasion probability and the magnitude of effects of successful mutants are also strongly affected by linkage. Mutants tightly linked to the polymorphic site have a relatively high invasion probability, which depends relatively weakly on the migration rate. In addition, their (average) effects are much smaller than for loosely linked mutations, which may reduce the possibility of detection. For moderate to strong migration, the invasion probability declines by several orders of magnitude as *r* increases from small values (say ≤ 0.01) to large values (close to 0.5). The waiting time for the establishment of such a mutation will be inversely proportional to its invasion probability. Furthermore, with increasing *r*, the mean effect of successfully invading mutations increases substantially (up to about four fold; see Fig. 6A). Thus, beneficial mutants will become established in the vicinity of locus B on a relatively short time scale, but will have predominantly small effects. By contrast, establishment of loosely linked or unlinked beneficial mutants will occur on a much longer time scale, but their effects will tend to be relatively large. These complex dependencies may severely affect the time scale on which clusters of locally beneficial mutations build up, as well as their detection.

## Acknowledgments

We thank Simon Aeschbacher for very helpful discussions, and Sabin Lessard and two anonymous reviewers for thoughtful comments on an earlier version which led to notable improvements. Financial support by the Austrian Science Fund (FWF) through the Vienna Graduate School of PopulationGenetics (GrantDK W1225-B20) to MP is gratefully acknowledged.

## Supporting information

The supporting information files *S*1 − *S*6 are Mathematica notebooks that can be found at http://phaidra.univie.ac.at/o:1181315.

## Appendix A The equilibrium *E*_*A*_ is externally unstable if it exists

If *m < a/*(1 + *a*), then there exists the equilibrium *E*_*A*_ at which *B*_2_ is fixed and *A*_1_ and *A*_2_ are in migration-selection balance. Due to symmetry with *E*_*B*_, we can derive the leading eigenvalue *λ*_*A*_ of the Jacobian by simultaneously changing *a* → *b* and *b* → *a* in equation (11). For the long-term dynamics the external stability (*λ*_*A*_ < 1) of this equilibrium is of interest. After some algebra this can be written as

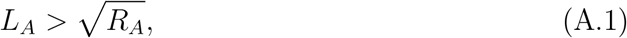

where *L*_*A*_ and *R*_*A*_ can be derived from the analogous expressions in the main text by using the same symmetry argument as above. Similarly, we can deduce 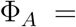 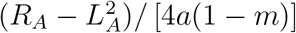. But here, the similarities end. It can be shown that

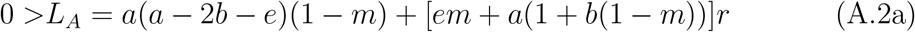

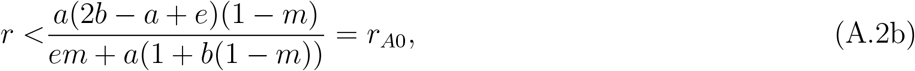

since [*em* + *a*(1 + *b*(1 − *m*))]*r* > 0 under our assumptions. Further, *r*_*A*0_ > 0, because also the numerator is positive. Thus, *E*_*A*_ is unstable if *r < r*_*A*0_.

Similarly to the proof of Proposition 1, we transform Φ_*A*_(*m*) to ψ_*A*_(*z*) = *d*_0_ + *d*_1_*z*, using *m*_*A*_ = *a/*(1 + *a*) as upper bound. Here,

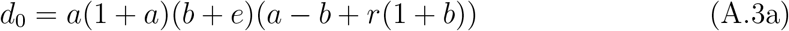

and

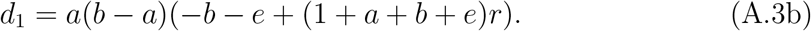

If *r* > (*b*+*e*)/(1+*a*+*b*+*e*) = *r*_*A*1_, then *d*_1_ > 0 and also *d*_2_ > 0, since (*b*+*e*)/(1+*a*+*b*+*e*) > (*b* − *a*)/(1 + *b*). Thus, if *r > r*_*A*1_, then Φ_*A*_(*m*) > 0, which implies *λ*_*A*_ > 1. However, it can be shown that *r*_*A*0_ *> r*_*A*1_.

In summary, we showed that *λ*_*A*_ > 1 for 0 < *r* < 1/2 if *E*_*A*_ is admissible and the same assumptions hold as in the analysis of *E*_*B*_.

## B Approximations of the invasion probability *π*_*i*_(*ϵ*) in (26)

Here, we derive the expression *β*(*ϵ*) given in (27), which plays a key role in the approximation of the invasion probability *π*_*i*_(*ϵ*) in (26), derived for a supercritical branching process by Athreya (1993) (see also Haccou et al., 2005, pp. 126-128). In addition, we show that the expression *β*_•_(*ϵ*) given by Aeschbacher and Bürger (2014) (eq. (64) in their Supporting Information, File S1) satisfies (29) and *β*_•_(*ϵ*) ≤ *β*(*ϵ*). However, if *r* > 0, in general the difference between *β*_•_(*ϵ*) and *β*(*ϵ*) is of order *O*(*ϵ*), so that *β*_•_(*ϵ*) does not always yield a first-order approximation in *E* of *π*_*i*_(*ϵ*). Nevertheless, for small values of *r*, *β*_•_(*ϵ*) often yields a more accurate approximation of *π*_*i*_(*ϵ*) than *β*(*ϵ*). In particular, in contrast to *β*(*ϵ*), it has the feature that the invasion probability may be maximized at a positive *r*.

We assume independent Poisson offspring distributions. Our starting point is the expression (5.81) in (Haccou et al., 2005, pp. 127), which in our notation from Section 4 reads

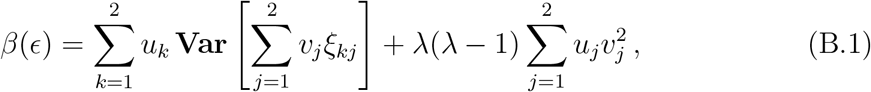

where *ξ*_*kj*_ are the Poisson variates for the offspring distribution, so that the expected number of offspring of type *j* of a type *k* parent is **E**[*ξ*_*kj*_] = *λ*_*kl*_. Therefore, **Var**[*ξ*_*kj*_] = *λ*_*kl*_. For notational simplicity, we omit the dependence of *λ*, *u*, and *v* on *ϵ*. Recall from Section 4 that the entries of the mean matrix **M** are the *λ*_*kl*_, and that *λ* is its principal eigenvalues.

We observe that

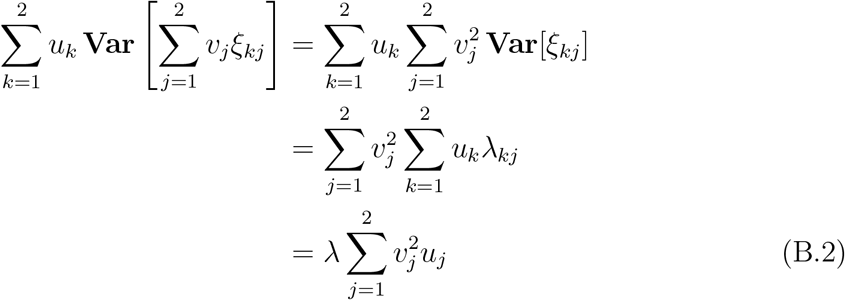

because *u* is the left principal eigenvector of **M**. Substituting (B.2) into (B.1) yields (27).

Aeschbacher and Bürger (2014, eq. (64) in File S1 of their Supporting Information) had obtained an erroneous expression (because of missing the square of *v*_*j*_ in the first equation of the derivation of B.2) which, after using *u***M** = *λu*, can be written as in (29). To prove *β*_•_(*ϵ*) ≤ *β*(*ϵ*), it is sufficient to show 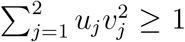. From the normalization (25), we obtain *u*_2_ = 1 − *u*_1_ and *v*_2_ = (1 − *u*_1_*v*_1_)/(1 − *u*_1_). Therefore,

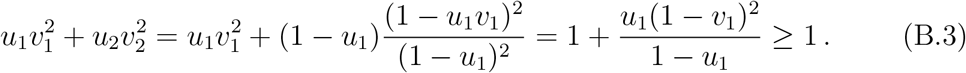

It is easily shown that 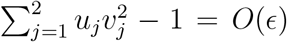, but not necessarily of order *O*(*ϵ*^2^). Therefore, *β*(*ϵ*) and *β*_•_(*ϵ*) may differ to first order in *ϵ*. Therefore, (26) shows that 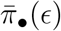 does not necessarily provide an approximation of *π*_*i*_(*ϵ*) to order *ϵ*.

However, if *r* = 0, *r* = *r*\*, *m* = 0, or *m* = *m*\*, we have *π*_•_(*ϵ*) = *π*(*ϵ*). Indeed, if *r* = 0, then **M** is a diagonal matrix with principal eigenvalue *λ* = 1 +(*a* + *e*)/(1 + *b*) and corresponding left and right eigenvectors *u* = (1, 0) and *v* = (1, 0)^*T*^. If *m* = 0, then **M** is a triagonal matrix with principal eigenvalue *λ* = 1 +(*a* + *e*)/(1 + *b*) and corresponding eigenvectors *u* = (1, 0) and *v* = (1, (1 + *a*)*r/*(*b* + *e* + (1 + *a*)*r*))^*T*^. In both cases (29) shows that

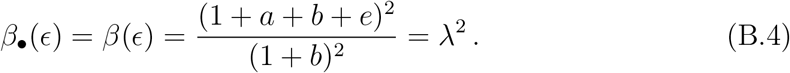

If *β*_•_(*ϵ*) coincides with *β*(*ϵ*), then so does the induced invasion probability. Finally, if *r* = *r*\* or *m* = *m*\*, then *ϵ* = 0, *λ* = 1 and *π*_*i*_(0) = 0 by (26).

Figures 2 and S2 compare the approximations 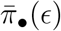 and 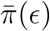 with the true mean fixation probability 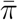 and highlight their accuracy.

## C Explicit slightly supercritical approximations for small *m* or small *r*

Because *λ*(*ϵ*) = 1 + *ρ*(*ϵ*) = 1 if *r* = *r*\* or *m* = *m*\*, we have *π*_1_(*ϵ*) = *π*_2_(*ϵ*) = 0 in these cases. In addition, for small *r* and based on *β*(*ϵ*), we obtain the approximation

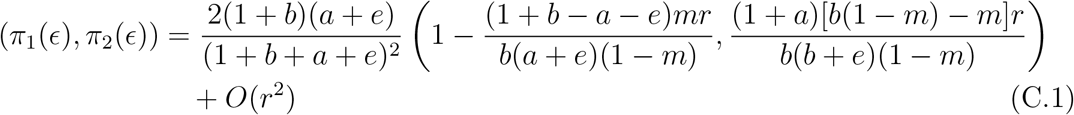

(File S3, Sect. 6). From (6), (7) and (C.1) we obtain for the mean invasion probability close to *r* = 0:

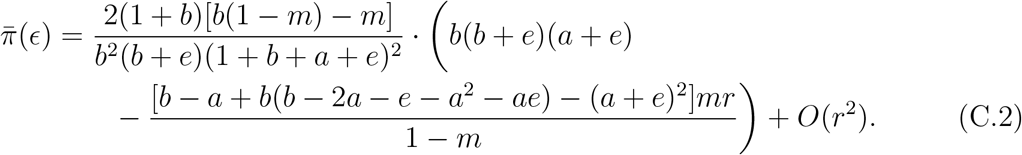

The leading-order term can be rewritten as in (31).

For small *m*, the first-order approximation of 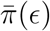 is more complicated, so we give only the leading term:

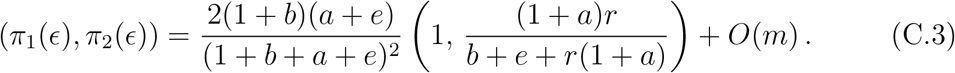

For 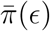, we obtained the following first-order approximation in *m* near *m* = 0:

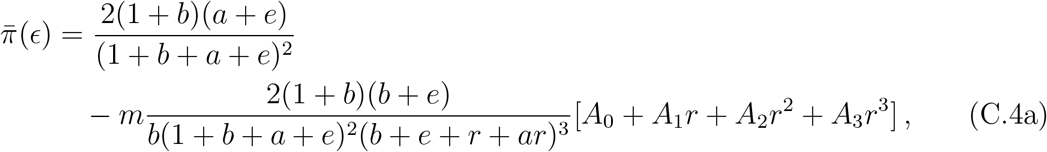

where

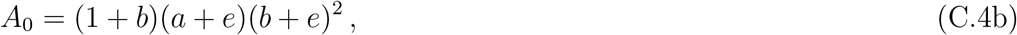

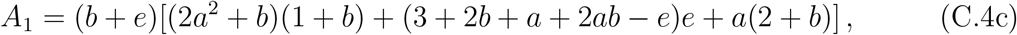

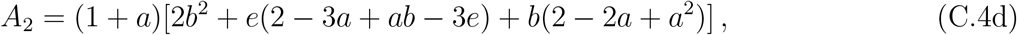

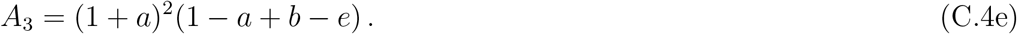

In the File S3, Sect. 4, we show that 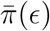 is decreasing for small *m*. We do this by showing that *A*_0_ + *A*_1_*r* + *A*_2_*r*^2^ + *A*_3_*r*^3^ > 0, if 0 < *r* < 1/2.

## D Explicit slightly supercritical approximations for weak evolutionary forces

For weak evolutionary forces (Section 3.2), the approximations for *π*_1_, *π*_2_, and 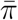 simplify (see File S5, Sect. 3):

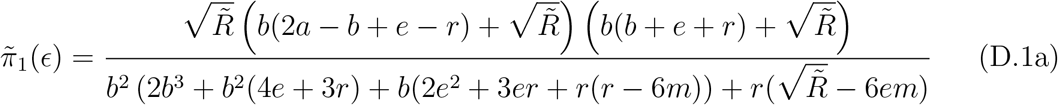

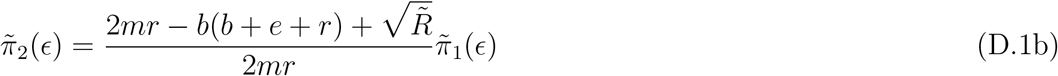

or shorter:

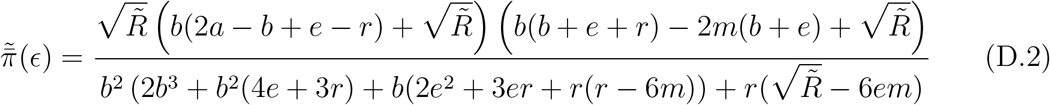

with 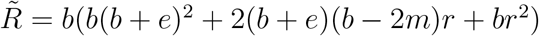.

## E Exact expressions and approximations for *r* = 0 and for *m* = 0

First, we derive an explicit solution for the average invasion probability if *m* or *r* is equal to zero. Second, we use this to derive a condition for the existence of a non-zero optimal recombination rate.

Because *π*_*i*_ = 1 − *σ*_*i*_, where (*σ*_1_, *σ*_2_) is the uniquely determined solution of (24) satisfying 0 ≤ *s*_1_ < 1 and 0 ≤ *s*_2_ < 1, we will differentiate the equations in (24) with respect to *r* and evaluate at *r* = 0. We write *s*_*i*_(0) for the values of *s*_*i*_ evaluated at *r* = 0. We note that if *r* = 0 or *m* = 0, then *λ*_12_ = 0 and 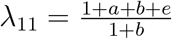. For simplicity, we set 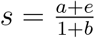 and write 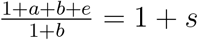, where *s* > 0.

Then *σ*_1_(0) satisfies

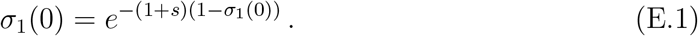

The solution can be written in terms of the product logarithm, or the Lambert function, i.e.,

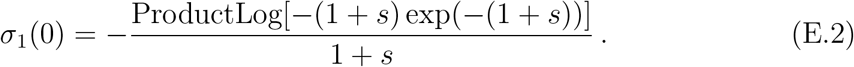

This has the series expansion

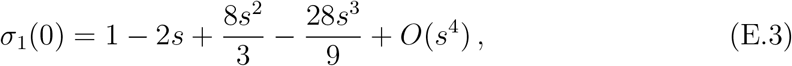

which implies

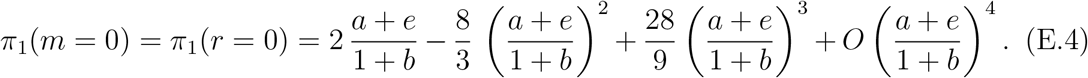

Because we assume *a < b*, we have *λ*_21_ < 1, *λ*_22_ < 1, and thus *σ*_2_(0) = 1.

Therefore, by (7) and (6), the explicit invasion probabilities are given to first order in 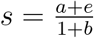 by

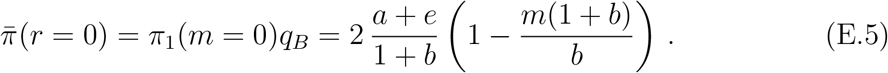

Higher-order approximations follows immediately from (E.4) and are very accurate.

If 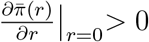, then there is a recombination rate that maximizes 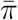 at a value greater than 0. By (7) we need the derivatives of *π*_1_ and *π*_2_ with respect to *r* at *r* = 0. We write *s*′_*i*_(0) for the values of the derivative of *s*_*i*_ with respect to *r* evaluated at *r* = 0.

Differentiation of (24) with respect to *r* and evaluation at *r* = 0 yields

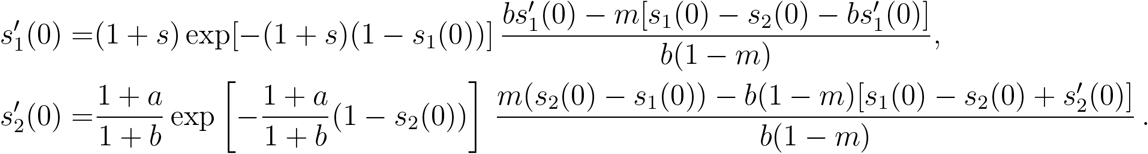

With *s*_*i*_ = *σ*_*i*_ and applying (E.1) and *σ*_2_(0) = 1, we obtain

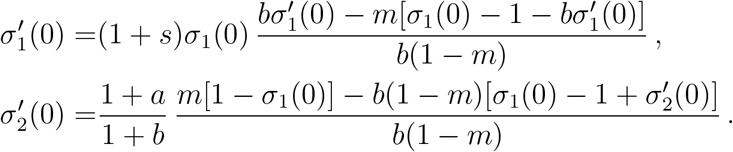

This is a linear system in *σ*′_1_(0) and *σ*′_2_(0), which has the following solution:

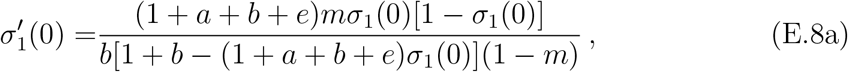

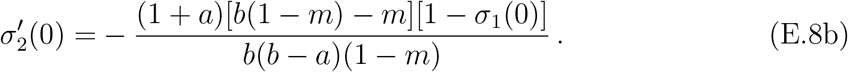

From (7), we obtain

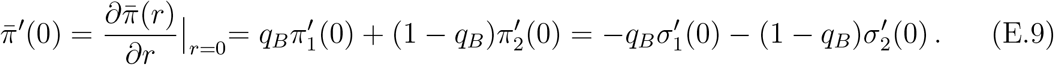

By substituting (E.8) and rearranging, we find

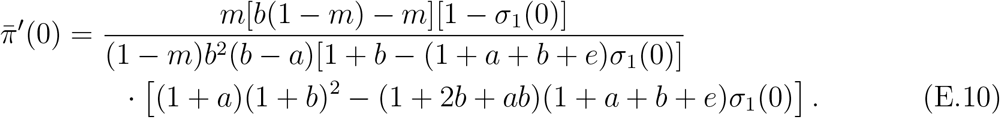

Therefore, the sign of the derivative at *r* = 0 is determined by

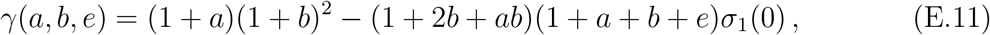

where *σ*_1_(0) is given by (E.2) with *s* = (*a* + *e*)/(1 + *b*). Because on the right hand side of (E.10) all factors other than *γ*(*a, b, e*) are positive, 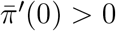 if and only if *γ*(*a, b, e*) > 0. In addition, by differentiation and using the monotonicity properties of the product logarithm, it can be shown that *γ*(*a, b, e*) is strictly monotone increasing in *a*, and there exists a unique value *a*\* > 0 for which *γ*(*a*\*, *b, e*) = 0 if −*a < e < b* − *x*, where *x* > 0 is of order *b*^2^.

By substituting *e* → *e*_*b*_*b* in *γ*(*a, b, e*) and performing a series expansion of *γ*(*a, b, e*_*b*_*b*) up to third order in *b* (assuming that *b* is small and |*e*_*b*_| ≤ 1), and finally resubstituting *e*_*b*_ by *e/b*, we find the following approximation for the critical value *a*\*:

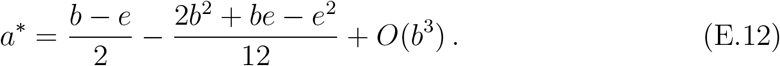

## SI Figures

**Figure S1:**
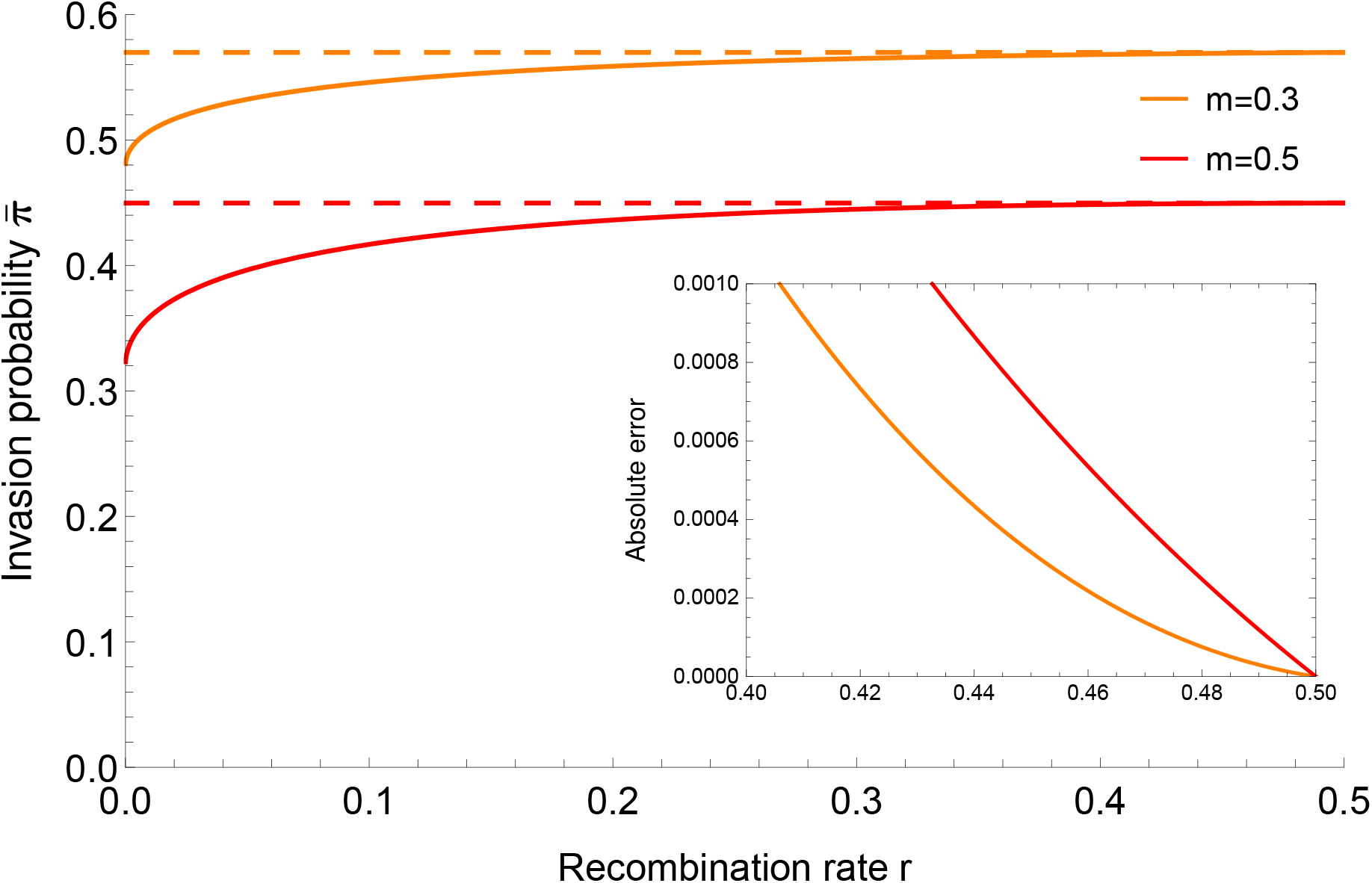
The invasion probability (solid curve) is shown as a function of the recombination rate. For both parameters combinations it is strictly increasing. Therefore, the optimum recombination rate is *r* = 0.5. The dashed lines show the constant values (~ 0.57 and ~ 0.45) of the maximum invasion probability achieved at *r* = 0.5. The other parameters values are *a* = 0.09, *b* = 0.1 and *e* = 0.75. The inset shows the difference between the optima and the invasion probability for large *r*.

**Figure S2:**
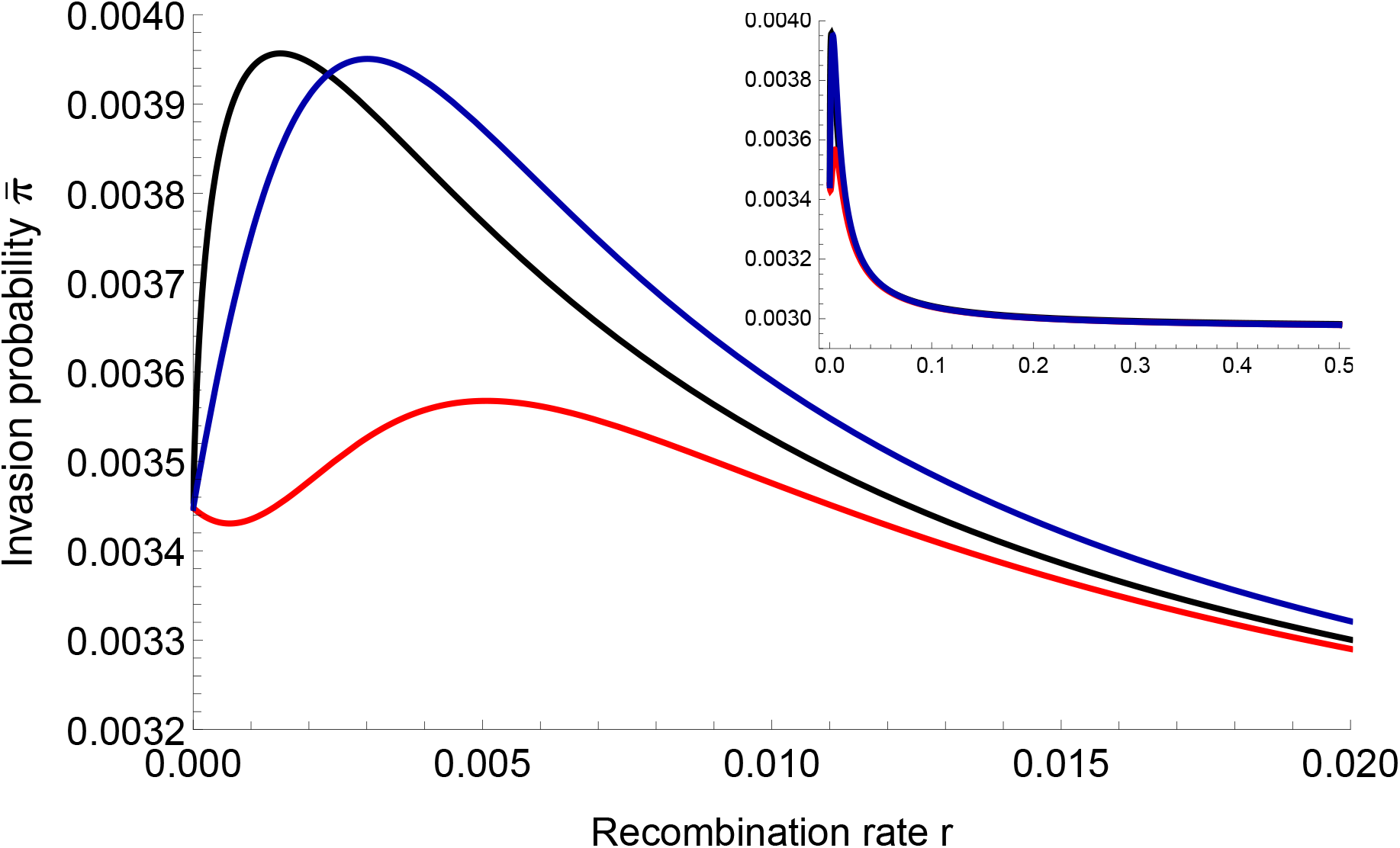
In this figure, the approximation 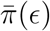 (red curve) has two local optima, one at *r* = 0 and another one for *r* > 0. The black curve is the numerical solution, whereas the blue curves correspond to 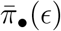. This figure shows that the slightly supercritical approximation 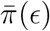 can behave quite differently than 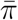 if linkage is very tight. The parameter values are *a* = 0.0035, *b* = 0.004, *e* = 0, and *m* = 0.002. Thus, *a*\* = 0.002. The inset shows the same plot for the full range of *r*.

**Figure S3:**
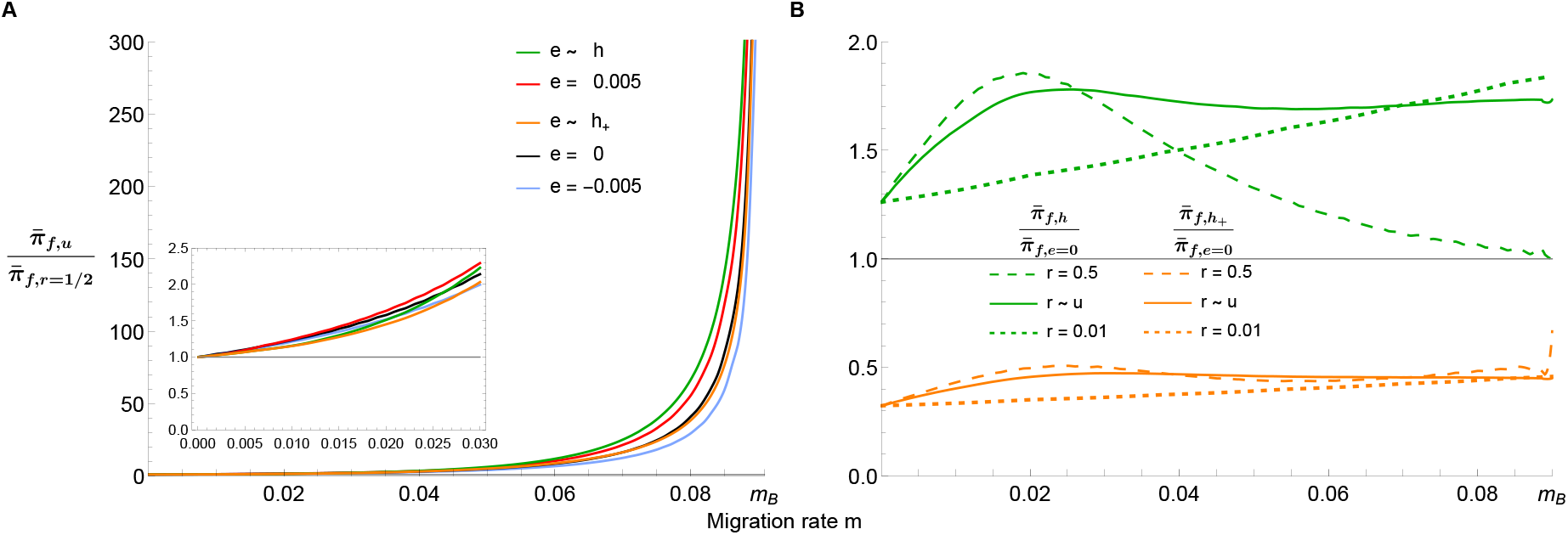
Ratios of averaged invasion probabilities of new mutations as functions of *m*. In all panels the invasion probability is averaged with respect to *f* (*a*) with mean 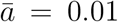. The only constant parameter is *b* = 0.1, so that the maximum possible migration rate is *m*_*B*_ ≈ 0.091. The purpose of this figure is to compare the effects of averaging over *r* with free recombination (panel A), and of averaging over *e* with *e* = 0 (panel B). For the epistatic values in Fig. 5A and the epistasis distributions in Fig. 5B, panel A displays the ratios of the invasion probabilities averaged over *a*, *r*, and potentially *e* (i.e., 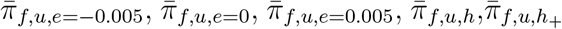), and the corresponding invasion probabilities for unlinked loci (e.g., 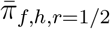). The line colors in A match those in Figs. 5A,B. Thus, the invasion probabilities shown by the solid curves in panels A and B are divided by those of the long-dashed curves of the same color. The curves tend to infinity at the value *m* at which the dashed curves in Fig. 5A and Fig. 5B reach zero. Panel B shows the ratios of the invasion probabilities averaged over *a* and *e* and of those without epistasis. The solid curves are obtained by additional averaging over *r*, i.e., they show 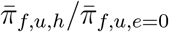 and 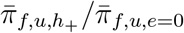. Thus, the invasion probability of each of the colored curves in Fig. 5B is divided by the invasion probability of the black curve in Fig. 5B of the same dashing style. The gray horizontal line in A and B is at 1 to provide a reference for the ratios. Roughly, these figures shows that tighter average linkage always has a positive effect (ratio > 1) on the invasion probability, whereas the direction of the effect of epistasis depends on the specific distribution.

**Figure S4:**
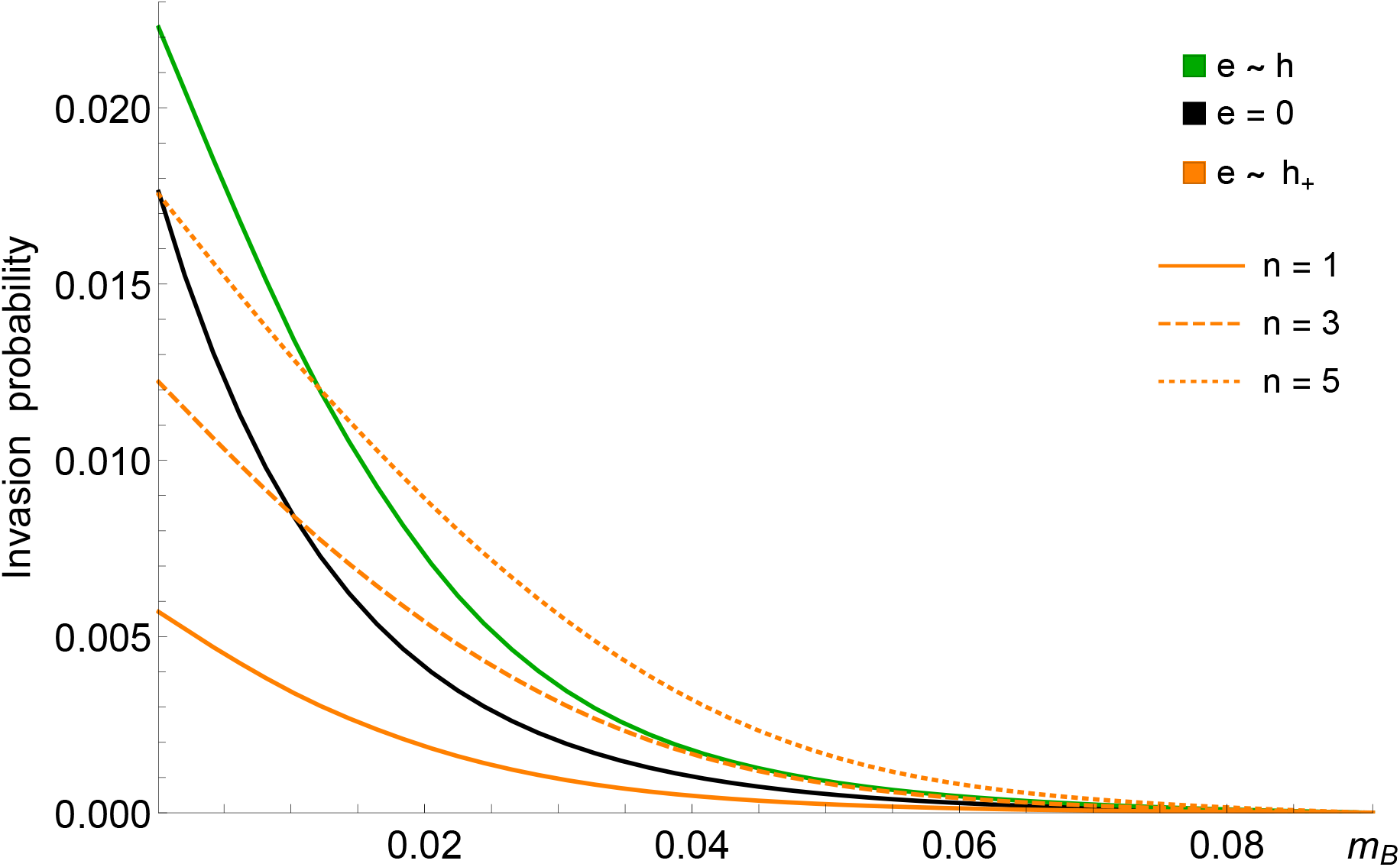
Averaged invasion probabilities as functions of *m*. The solid curves in this figure are the same as in Fig. 5B. In all cases, the additive effects *a* follow the exponential distribution *f* with 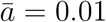, the recombination probabilities *r* are uniformly distributed on 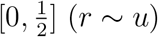, and *b* = 0.1. The black curve shows the invasion probability averaged over *a* and *r* with *e* = 0, i.e., 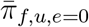. For the other curves, *e* is drawn from a normal distribution. The green curve shows 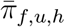, the solid orange curve shows 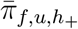 with *n* = 1 in (47). The long-dashed and short-dashed solid orange curve show 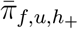 with *n* = 3 and *n* = 5, respectively, in (47). Comparison of the orange curves demonstrates that for the distribution *h*_+_ a higher variance of the epistatic effects increases the invasion probability. For large enough *m*, the resulting invasion probability can even exceed that caused by the distribution *h*, which has mean 0.

**Figure S5:**
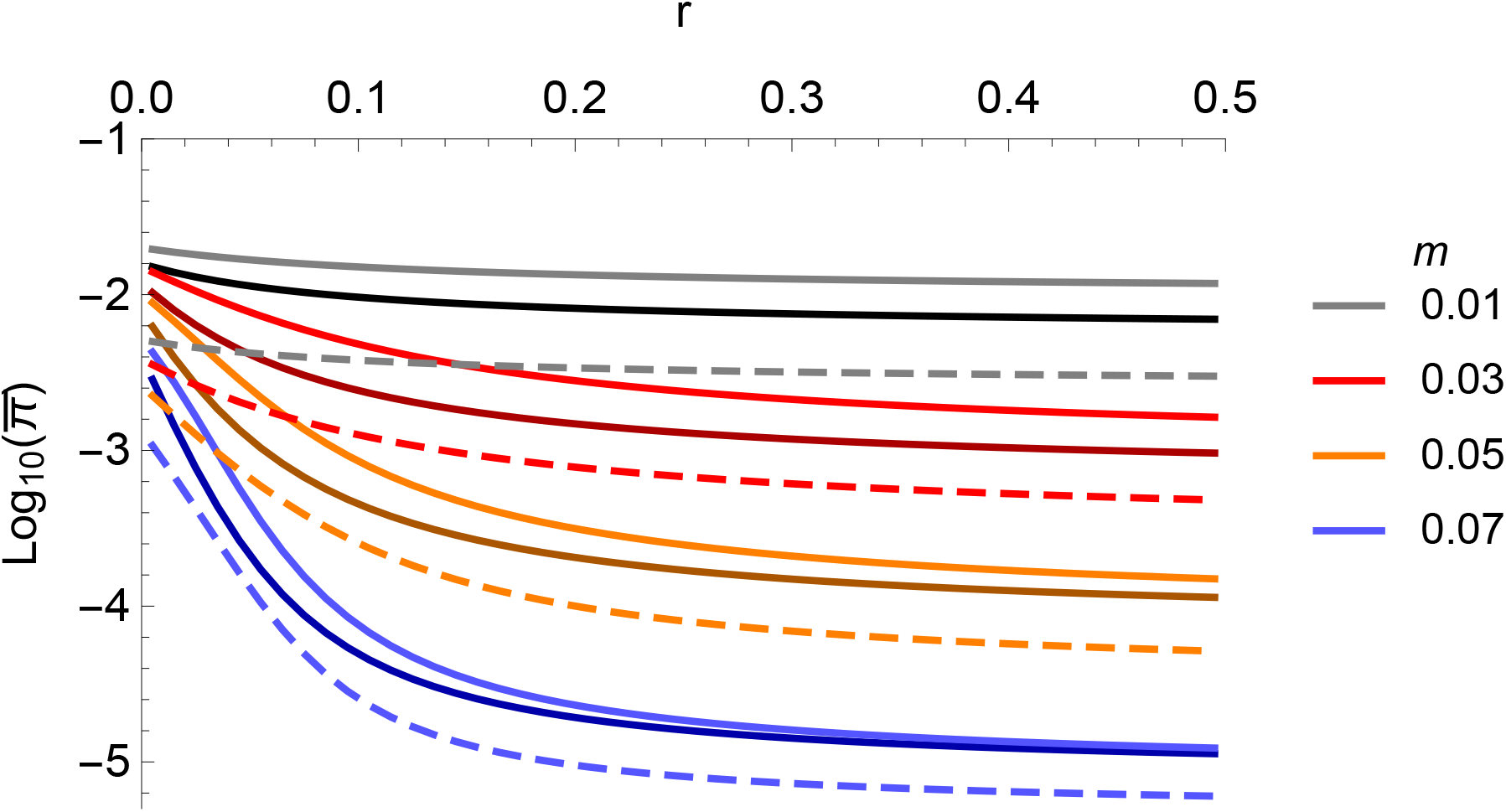
Averaged invasion probabilities corresponding to Fig. 6 as functions of the recombination rate on a logarithmic scale. Shown are 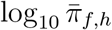 (bright colors, solid), 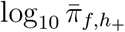 (bright colors, dashed), and 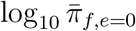 (dark colors, solid). The log10 scale is chosen for improved visibility. The line colors indicate the migration rate (legend).

**Figure S6:**
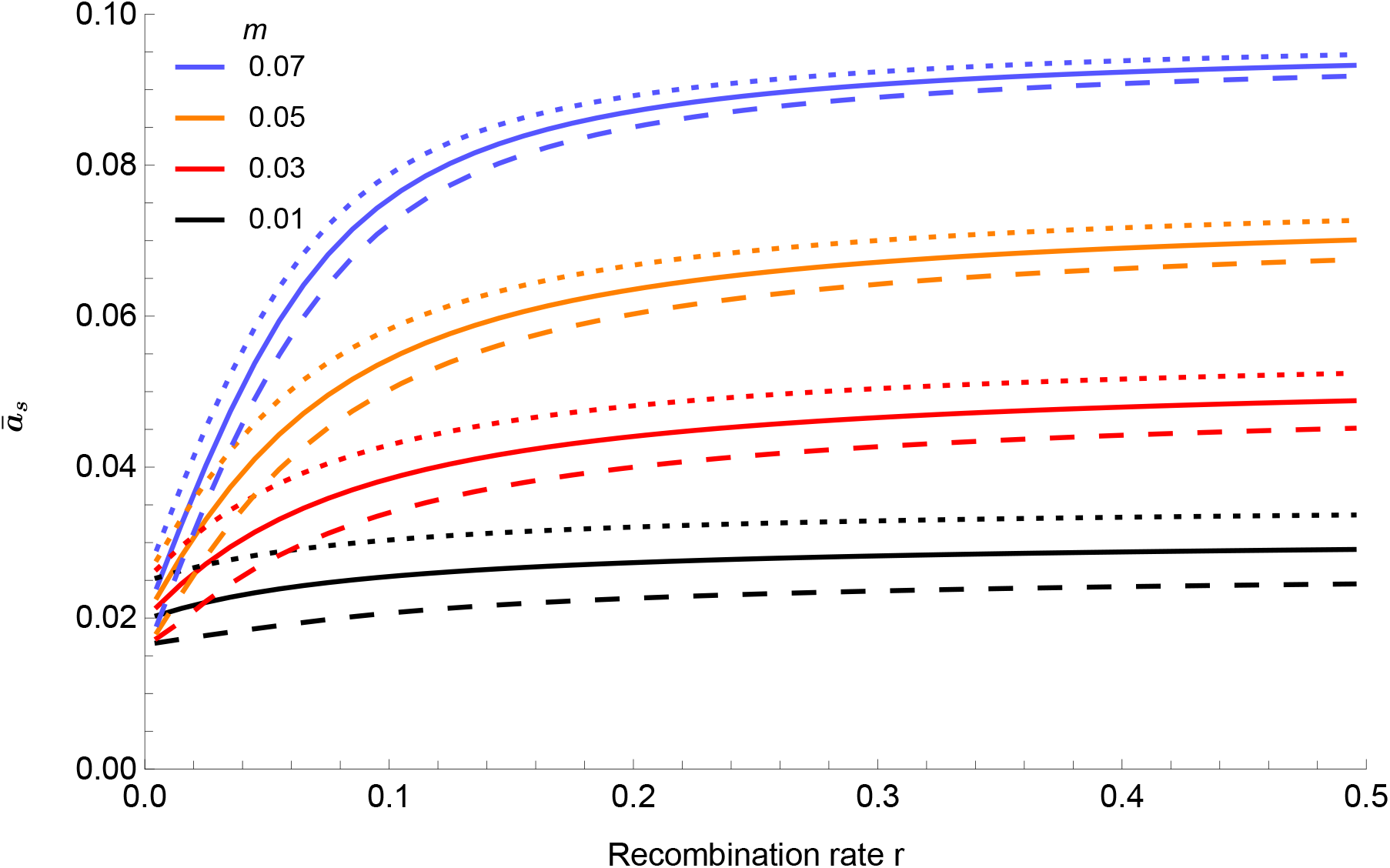
This figure is analogous to Fig. 6A, except that here epistatic values are not drawn from a distribution. It shows the average value 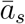 of successfully invading mutations for three fixed values of *e*. Additive effects are drawn from an exponential distribution with mean 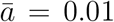. Solid curves are for *e* = 0. Dotted curves are for *e* = −0.005 and dashed curves for *e* = 0.005. We observe that smaller values of *e* lead to, on average, larger additive fitness values 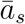 of the successful mutations.

**Figure S7:**
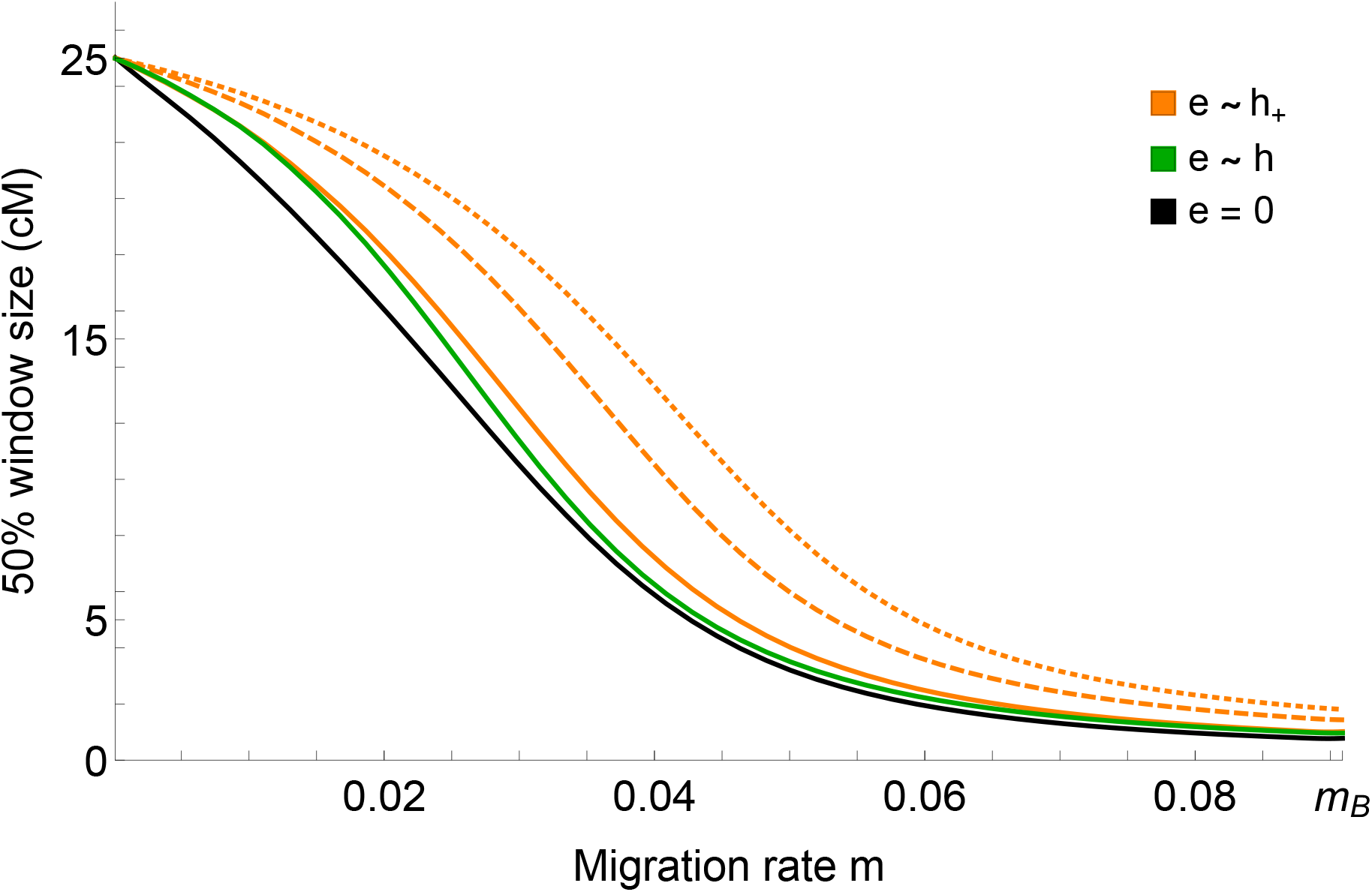
This is analogous to Fig. 8. Here, we include *C*_50_ for the distribution *h*_+_ with three values of *n* in (47), where *n* is the dimension in Fisher’s geometric model (Blanquart et al., 2014). The solid orange curve is for *n* = 1 (as in Fig. 8), the dashed orange curve is for *n* = 3, and the dotted orange curve is for *n* = 5. The corresponding variances are 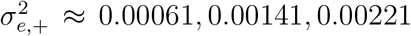, respectively. Therefore, about 10.2%, 20.2%, and 25.2% of the epistatic effects are positive. Thus, the higher the variance of a normal distribution of *e* with negative mean (*h*_+_), the higher is the increase in window size. In particular, the window size can be substantially increased relative to no epistasis (*e* = 0) or to normally distributed epistatic values with mean 0 and small variance (as for *h*).

